# Comprehensive IsomiR sequencing profile of human pancreatic islets and EndoC-βH1 beta-cells

**DOI:** 10.1101/2023.11.08.566223

**Authors:** Stefano Auddino, Elena Aiello, Mattia Toniolli, Giuseppina Emanuela Grieco, Daniela Fignani, Giada Licata, Marco Bruttini, Alessia Mori, Andrea Berteramo, Erika Pedace, Laura Nigi, Caterina Formichi, Giuseppe Quero, Vincenzo Tondolo, Gianfranco Di Giuseppe, Laura Soldovieri, Andrea Mari, Andrea Giaccari, Teresa Mezza, Francesco Dotta, Guido Sebastiani

**Author notes:** **Corresponding Author** Prof. Francesco Dotta, Diabetes Unit, Dept. of Medicine, Surgery and Neurosciences, University of Siena, Siena, Italy. Shared First co-authorship (Stefano Auddino, Elena Aiello).

## Abstract

**Aims/Hypothesis:** MiRNAs play a crucial role in regulating the islet transcriptome, influencing beta cell functions and pathways. Emerging evidence suggests that during biogenesis a single miRNA locus can generate various sequences, known as isomiRNAs (isomiRs). However, a comprehensive profiling analysis of isomiRs in human pancreatic islets and beta cells is still lacking. This study aims to unveil the isomiRs expression profile in Laser Capture Microdissected (LCM) human pancreatic islets (HI) from non-diabetic donors and in the human beta cell line EndoC-βH1, in order to shed light on novel molecular mechanisms governing beta cell function.

**Methods:** RNA was extracted from LCM HI deriving from n=19 non-diabetic donors and from EndoC-βH1 beta cells. Small RNA-seq was performed. Data were processed with the sRNAbench online pipeline for miRNAs/isomiRs quantification. Results were further validated using an external miRNA-seq database (isomiRdb).

**Results:** In both HI and EndoC-βH1, isomiRs accounted for a substantial proportion of total miRNA reads (HI: 59.4+/-1.9%; EndoC-βH1: 43.8+/-0.6%). Among isomiRs, the most prevalent types were 3’-end modifications, including trimming (HI=71.8+/-2.8%; EndoC-βH1=55.8+/-1.0%) and extension (HI: 12.1+/-1.9%; EndoC-βH1: 17.4+/-0.9%), followed by non-templated addition (HI: 9.8+/-0.9%; EndoC-βH1: 14.0+/-1.2%). The analysis of the composition of the n=10 most expressed miRNAs highlighted a significant contribution of reads assigned to isomiRs. For instance, the most abundant miRNA, miR-375-3p, resulted from 59.7+/-2.4% of canonical and 40.3+/-2.4% of isomiRs in EndoC-βH1 and from 45.3+/-2.0% of canonical and 54.7+/-2.0% of isomiR reads in HI. Interestingly, miR-7-5p, a beta cell-specific miRNA, was predominantly expressed as an isomiR both in EndoC-βH1 (65.3+/-2.7%) and in HI (82.4+/-1.4%). To identify a reliable beta cell isomiR signature, common sequences detected in HI and EndoC-βH1 were filtered based on their contribution to total miRNA expression, ultimately resulting in a set of 46 isomiRs. The expression of the isomiR signature in beta cells was further evaluated using an external database, isomiRdb, which contains small-RNA sequencing data from 99 different human cell types. This analysis revealed a significant enrichment of 11 out of the 46 isomiRs in beta cells compared to other cell types. The signature was functionally characterized through regression analysis with clinical and metabolic parameters related to beta cell function in non-diabetic individuals, demonstrating a significant negative correlation between basal insulin secretion and isomiR-411-5p, but not with its corresponding canonical miRNA.

**Conclusion/Interpretation:** This study provides a comprehensive profile of isomiR expression in pancreatic islets and beta cells, highlighting the potential significance of isomiRs as novel regulators of beta cell function.

**Graphical Abstract:** 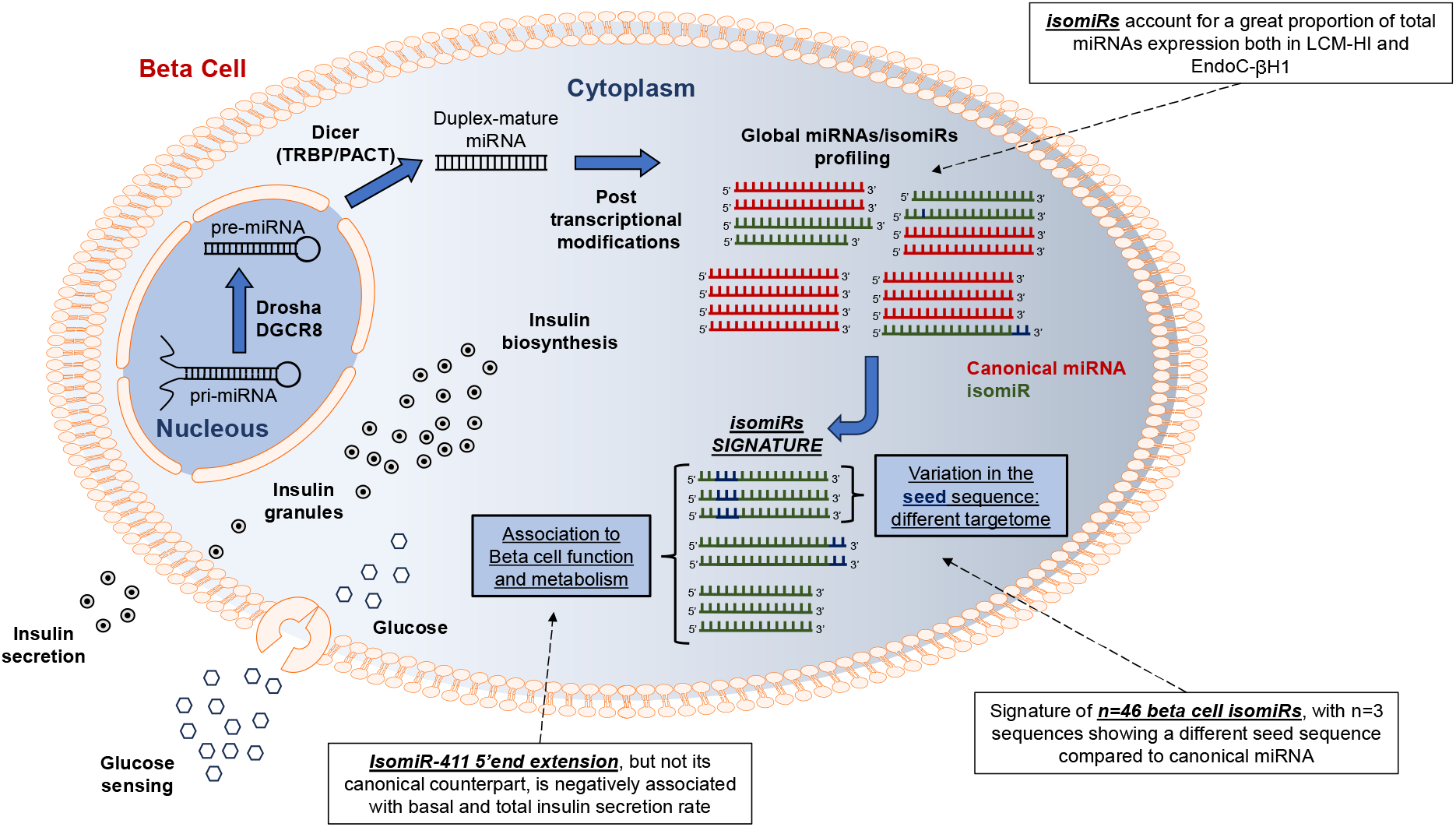

**Research in Context:** *What is already know about this subject?:*

- isomiRs are sequence variants of microRNAs (miRNAs) and may have distinct functional role respect to the canonical sequence.
- isomiRs show cell/tissue specificity and are involved in multiple disease contexts.

*What is the key question?:*

- What is the profile of isomiRs in human pancreatic islets (HI) and in beta cells?
- Do isomiRs have a functional role in beta cells?

*What are the new findings?:*

- IsomiRs represent a relevant fraction of total miRNAs in HI and beta cells.
- 3’end miRNA sequence modifications are the major fraction of isomiRs in beta cells.
- A set of n=11 isomiRs, more expressed than their canonical miRNAs, are enriched in beta cells compared to the other human cell types.
- Specific isomiRs are associated with beta cell glucose sensitivity and basal insulin secretion.

*How might this impact on clinical practice in the foreseeable future?:*

- A comprehensive profile of isomiRs in beta cells may improve our understanding of molecular mechanisms driving beta cell function and dysfunction.
- A highly detailed and granular view of miRNAs sequence variants and their expression levels may help in the design of novel therapeutic RNA-based strategies.

## Introduction

MicroRNAs (miRNAs) are ∼22 nucleotides-long non-coding RNA transcripts involved in the regulation of genes expression by pairing to messenger RNA (mRNA) target sequences [1–3]. The major determinant of miRNAs binding to target mRNAs is a 6-8 nucleotides domain at the 5’ end of miRNA (*seed* sequence). The perfect pairing between miRNA *seed* and the corresponding mRNA target sequence most likely results in the reduction of its expression. Hence, the targetome of each miRNA is largely determined by *seed* sequence. Consequently, in light of their major role on genes expression modulation, miRNAs regulate pivotal biological processes [2], and many of them are altered in multiple diseases, being addressed also as therapeutic targets [4].

Known and mapped miRNAs are listed in public databases, such as miRbase (www.miRbase.org) [5]. MiRNAs included in miRbase have been defined as a single sequence of RNA (i.e. ‘canonical’ miRNA). However, next generation sequencing (NGS) studies performed in multiple biological contexts revealed a relevant proportion of miRNA sequences which differed in length and/or one or more nucleotides from canonical miRNAs [6–8]. MiRNAs sequence variants, called isomiRNAs or isomiRs, were initially dismissed as sequencing errors or artifacts. Nevertheless, the refinement of sequencing technologies and bioinformatic analytical pipelines confirmed the bone-fide of these miRNA sequence variants [9, 10].

The biogenesis of isomiRs is strictly related to the maturation and/or editing of canonical miRNAs, and involves alternative miRNA precursor (pre-miRNA) cropping and/or dicing, terminal nucleotide trimming or addition (5’ or 3’ trimming/extension), non-templated nucleotides addition (NTA) and single nucleotides modification due to RNA editing, all operated by specialized enzymes. However, the exact molecular mechanism of their biogenesis remains unclear. Nonetheless, isomiRs have been shown to be tissue-specific [11] and, for example, used to classify multiple cancer-types [12]. Of importance, isomiRs can be functionally associated with RISC complex, thus able to inhibit mRNA expression [13, 14]. The function of isomiRs may vary respect to the canonical ones depending on the sequence variation type, thus broadening the role of a specific miRNA or having a completely independent role in regulating physiological or pathological processes. Particularly, variations at the 5’ end of canonical miRNAs (both trimming and extensions) can modify the seed sequence thus changing its targetome; alternatively, trimming or extension at 3’ end of canonical miRNAs can modify its stability/half-life, Argonoute-2 loading or non-canonical target recognition, thus significantly impacting on final function.

The advent of miRNAs high-throughput analytical platforms bolstered the characterization of their expression profiles in multiple cells and tissues [15], revealing a landscape of ubiquitary and/or cell/tissue-specific miRNAs and isomiRs [16]. Multiple studies also described the profile of miRNAs in pancreatic islets and in beta cells, showing a peculiar fingerprint characterized by the high expression of specific miRNAs (i.e. miR-375, miR-7) alongside with others involved in the regulation of specialized endocrine islet cells functions [17–20]. Of note, additional studies further clarified the role of these miRNAs in beta cell functions, demonstrating their activity in development and differentiation [21–24], proliferation [25–29], apoptosis [30–36] and, importantly, basal- and/or glucose-stimulated insulin secretion [17, 37–44].

Given the growing diversity in both the sequence and functional roles of each miRNA, it is crucial to re-examine their expression patterns at the sequence level in human pancreatic islets (HI) and beta cells. This re-evaluation aims to deepen our understanding of the molecular mechanisms underlying miRNA regulation of beta cell functions and may shed light onto novel mechanisms which can be therapeutically treated. Although there have been previous indications of isomiRs playing a role in pancreatic islets [45], a comprehensive characterization of isomiRs, especially considering recent advancements in technological platforms and bioinformatic pipelines, has yet to be undertaken.

In this study, we profiled isomiRs in laser-capture microdissected (LCM) HI of non-diabetic living subjects and in the human beta cell line EndoC-βH1, with the aim of establishing a distinct signature of isomiRs specific to pancreatic islet and beta cells. We employed an up-to-date next-generation sequencing (NGS) protocol and applied a rigorous bioinformatic pipeline to analyze a robust and reliable set of isomiRs. Our findings revealed that a significant portion of relevant miRNAs, highly expressed in HI and beta cells, predominantly exist as isoforms, with distinct sequences and potentially alternative functions compared to the canonical forms. Additionally, through the validation in an external sequencing dataset, we identified a putative isomiRs signature specific to beta cells. This discovery represents the first instance of uncovering a putative beta cell-derived isomiR signature with defined role in this context.

## Methods

### Study subjects and human pancreas tissue collection

Human pancreatic sections analyzed in this study were obtained from pancreata of n=19 donors. Specifically, n=3 brain-dead adult non-diabetic multiorgan donors collected in the European Network for Pancreatic Organ Donors with Diabetes (EUnPOD-INNODIA consortium), and n=16 non-diabetic living donors undergoing pylorus-preserving pancreatoduodenectomy recruited at the Digestive Surgery Unit and studied at the Centre for Endocrine and Metabolic Diseases Unit (Agostino Gemelli University Hospital, Rome, Italy) Indications for surgery were periampullary tumors, pancreatic intraductal papillary tumors, mucinous cystic neoplasm of the pancreas, and nonfunctional pancreatic neuroendocrine tumors (**ESM Table 1**). Pancreatic tissue specimens were immediately snap frozen in liquid nitrogen and stored at −80°C embedded in Tissue-Tek OCT compound.

**Table 1.** IsomiR type fractions in human pancreatic islets and EndoC-βH1. The table shows the contribution of the different isomiRs classes to the total isomiRs expression in LCM-HI samples and EndoC-βH1. The average contribution (reported as percentage of total isomiRs content) and the standard deviation across samples are represented.

| **IsomiR Type** | **Human islets**<br>**(% of total**<br>**isomiRs)** | **EndoC-βH1**<br>**(% of total**<br>**isomiRs)** |
| --- | --- | --- |
| Ext 3p | 12.06±1.86 | 17.4±0.88 |
| Ext 5p | 0.15±0.04 | 1.21±0.13 |
| MultiVariant | 0.73±0.14 | 5.93±0.6 |
| NTA | 9.8±0.88 | 13.99±1.22 |
| NucVar | 5.06±1.31 | 3.96±1.88 |
| Trim 3p | 71.79±2.76 | 55.79±0.95 |
| Trim 5p | 0.4±0.11 | 1.72±0.02 |

The study protocol (ClinicalTrials.gov NCT02175459) was approved by the Ethical Committee Fondazione Policlinico Universitario Agostino Gemelli IRCCS – Università Cattolica del Sacro Cuore (P/656/CE2010 and 22573/14), and all participants provided written informed consent, followed by a comprehensive medical evaluation, as previously described [46, 47]

In INNODIA EUnPOD network, pancreata not suitable for organ transplantation were obtained with informed written consent by organ donors’ next-of -kin and processed with the approval of the local ethics committee of the Pisa University (**ESM Table 2**).

**Table 2.** IsomiRs composition of the n=10 most expressed miRNAs in Human pancreatic islets. The table shows the isomiRs composition of the n=10 most expressed miRNAs in LCM-HI samples (n=19). The average proportion (percentage) and its standard deviation across samples are reported. MiRNAs are listed from the most expressed (i.e. let-7b-5p) to the least expressed (miR-148a-3p).

| miRNA | Ext 3p | NTA | Trim 3p | Canonical | NucVar |
| --- | --- | --- | --- | --- | --- |
| hsa-let-7b-5p | 23.7±2.2 | 7.1±0.53 | 32.8±3.1 | 36.2±1.4 | 0 |
| hsa-let-7a-5p | 1.42±0.2 | 0 | 43.0±2.3 | 51.3±2.15 | 4.1±1.1 |
| hsa-let-7f-5p | 3.45±0.5 | 1.4±0.13 | 44.0±3.7 | 49.6±3.81 | 1.4±0.4 |
| hsa-miR-375-3p | 0 | 15.4±1.2 | 36.0±3.2 | 45.2±2.0 | 3.2±0.9 |
| hsa-miR-7-5p | 0 | 0 | 82.4±1.3 | 17.5±1.3 | 0 |
| hsa-miR-125a-5p | 0 | 1.5±0.37 | 96.7±0.6 | 1.7±0.4 | 0 |
| hsa-let-7i-5p | 0 | 6.7±0.97 | 8.2±0.9 | 67.9±4.0 | 17.1±4.5 |
| hsa-miR-16-5p | 1.22±0.3 | 0 | 4.7±1.0 | 79.4±3.4 | 14.5±3.7 |
| hsa-let-7e-5p | 3.93±0.9 | 3.8±0.49 | 47.7±2.3 | 43.1±2.2 | 1.4±0.4 |
| hsa-miR-148a-3p | 2.02±0.4 | 11.9±1.27 | 13.6±2.0 | 57.6±3.2 | 14.7±4.2 |

Living donors were metabolically profiled before undergoing surgery and subjected to a 75-g oral glucose tolerance test and glycated haemoglobin (HbA1c) testing to exclude diabetes, according to the American Diabetes Association criteria. None of the patients enrolled had a family history of diabetes. For detailed methods regarding metabolic screening please refer to the ESM methods section.

### Laser Capture Microdissection

Pancreatic human tissues were cryosectioned and subjected to Laser Capture Microdissection (LCM) using an Arcturus XT Laser-Capture Microdissection system (Arcturus Engineering, Mountain View, CA, USA) by melting thermoplastic films mounted on transparent LCM caps (cat. LCM0214-ThermoFisher Scientific, Waltham, MA, USA) on specific islet areas. Intrinsic beta cells autofluorescence allowed the identification of human pancreatic islets for LCM procedure (see ESM Methods).

### EndoC-βH1 cell culture

EndoC-βH1 human beta cell line [48] was obtained by UniverCell-Biosolutions (Toulouse, France) and used for all experiments between passages 78-88. EndoC-βH1 were cultured at 37 °C with 5% CO_2_ in coated flask and maintained in culture in low-glucose DMEM (cat. D6046) (see ESM Methods).

### RNA extraction and quality control

Total RNA was extracted from each LCM sample using PicoPure RNA isolation kit Arcturus (cat. kit0204 - ThermoFisher Scientific, Waltham, MA, USA) following manufacturer’s procedure.

Total RNA was extracted from approximately 3.0 × 10^5^ EndoC-βH1 using Direct-zol RNA Miniprep Kit (cat. R202-Zymo Research, Irvine, CA, US) following manufacturer’s instructions.

In order to evaluate RNA abundance and purity, Agilent 2100 Bioanalyzer with RNA Pico chips (cat. 5067-1513 Agilent Technologies, Santa Clara, CA, USA) was performed for each RNA sample reporting RNA integrity (RIN) and concentration and by excluding samples with RIN<5.0 (**ESM Table 3**) (see ESM Methods).

**Table 3.** IsomiRs composition of the n=10 most expressed miRNAs in the human beta cell line EndoC-βH1. The table shows the isomiRs composition of the n=10 most expressed miRNAs in EndoC-βH1 samples (n=3). The average proportion (percentage) and its standard deviation across samples are reported. MiRNAs are listed from the most expressed (i.e. miR-375-3p) to the least expressed (miR-192-5p).

| **miRNA** | **NTA** | **NucVar** | **Trim 3p** | **Canonical** | **Ext 3p** | **MultiVariant** | **Trim 5p** |
| --- | --- | --- | --- | --- | --- | --- | --- |
| hsa-miR-375-3p | 9.5±0.7 | 1.4±0.4 | 29.2±2.0 | 59.6±2.4 | nd | nd | nd |
| hsa-miR-16-5p | nd | 6.2±3.2 | 2.1±0.3 | 89.6±2.4 | 2.0±0.4 | nd | nd |
| hsa-miR-7-5p | nd | nd | 54.1±1.9 | 34.7±2.2 | 11.1±0.2 | nd | nd |
| hsa-let-7a-5p | nd | nd | 26.1±0.16 | 71.2±0.1 | 2.5±0.09 | nd | nd |
| hsa-let-7f-5p | 1.2±0.1 | nd | 24.3±0.7 | 70.2±0.7 | 4.1±0.03 | nd | nd |
| hsa-let-7i-5p | 4.7±0.3 | 4.8±2.6 | 6.0±0.7 | 84.3±1.8 | nd | nd | nd |
| hsa-miR-432-5p | 27.9±3.3 | nd | 40.6±2.2 | 31.3±1.0 | nd | nd | nd |
| hsa-miR-183-5p | nd | nd | 2.1±0.4 | 52.2±0.6 | 15.2±1.0 | 13.7±1.0 | 16.5±1.0 |
| hsa-miR-200c-3p | 6.1±0.2 | nd | 55.1±0.7 | 38.6±0.6 | nd | nd | nd |
| hsa-miR-192-5p | 16.2±0.9 | nd | 1.4±0.3 | 23.8±0.5 | 3.7±0.8 | 52.5±2.1 | 2.2±0.1 |

### Small RNA sequencing

Small RNA Seq was performed using QiaSeq miRNA library kit coupled with Illumina platforms sequencing. A total of 1 ng of RNA from microdissected human pancreatic islets and from EndoC-βH1 was used to generate cDNA libraries using QiaSeq miRNA library kit (Qiagen, Hilden, Germany) following manufacturer’s instructions (see ESM Methods).

### sRNAbench pipeline for small RNA-seq

FastQ files obtained from the sequencing were analyzed with the *sRNAbench* online pipeline [49] and processed with the Qiagen UMIs protocol for adapters and duplicates removal. To analyse isomiRs, a strong quality filter step was implemented to ensure a high quality of each base in the read. Reads that passes the quality filters were then aligned in genome mapping mode with the bowtie algorithm using the seed option and annotated to miRbase release 22.1 for miRNAs/isomiRs quantification (see ESM methods).

### IsomiR analysis and filtering steps

The raw counts matrix was obtained from the *microRNAannotation* file (*sRNAbench* output) of each sample. Several filtering steps were implemented to remove ambiguous sequences and potential sequencing errors from the analyses. In details, a relative abundance filtering, based on the contribution of the sequence to the total miRNA expression, was implemented to remove false isomiRs from the analysis. After the filtering steps, remaining sequences were assigned to the different isomiR types (or classes) according to the modification that distinguish them from the canonical miRNA (see ESM Methods).

### *In-silico* Sequencing simulation

To evaluate the goodness of the relative abundance filtering in removing possible sequencing errors from the analysis, an in-silico simulation was performed. A sequence of 22 nucleotides was generated, thus representing a canonical miRNA. The probability of the sequencing error was modelled considering the lowest quality score allowed in the analysis (Q=30). The sequencing of the generated read was simulated *n=1,000,000* times to model the probability of false isomiRs originating from sequencing errors (see ESM Methods).

### Beta cell isomiR validation analysis in isomiRdb

IsomiRdb datasets were accessed and re-analysed to validate the isomiR signature in beta cells and other cell types. Sample metadata, miRNA expression (normalized in Read Per Million), and isomiR expression (Reads Per Million) files were downloaded from isomiRdb, and re-analysed using a multi-step filtering (see ESM Methods).

### Statistical Analysis

Regression analyses between isomiRs expression and each clinical parameters were performed with linear regression models. In the models isomiR expression was used as dependent variable and the clinical parameter, the age, the gender, and the BMI as regressors. For each model influent points were detected and removed. Regressions with a p-value associated to the coefficient assigned to the clinical parameter < 0.05 were considered as statistically significant, independently from the effect of the covariates (see ESM Methods).

Hierarchical clustering analysis of miRNAs composition was performed independently for HI and EndoC-βH1 samples. The analysis was aimed to identify clusters of miRNAs with similar composition in the different isomiR classes for the two experimental groups. The similarity in miRNAs composition among clusters detected in HI and EndoC-βH1 was computed using the Jaccard similarity coefficient (see ESM Methods).

Delta Ranking analysis was performed to identify isomiR sequences with a higher contribution in expression compared to their corresponding canonical counterparts in each experimental group. Sequences were ranked in a descending way, and the difference in rank between each isomiR and its canonical sequence was computed. Highly negative values of delta rank are related to isomiR sequences much more expressed than the corresponding canonicals (see ESM Methods).

All the statistical and bioinformatic analyses were performed in R (version 4.2.2)[50].

## Results

### Implementation of the Sequencing Analysis Pipeline to Detect a Reliable IsomiR Signature in Human pancreatic Islets and Beta-Cells

IsomiRs are considered non-artifactual variants of miRNA sequences [9]. Consequently, due to their potential biological significance, a comprehensive characterization of their expression profiles across various tissutal and cellular contexts becomes of high importance. To analyze the abundance and distribution of isomiRs in human pancreatic islets (HI) and beta cells, we collected total RNA samples from LCM HI from n=3 non-diabetic multiorgan donors and n=16 normal glucose-tolerant (NGT) living subjects, and from n=3 different passages of the human beta cell line EndoC-βH1. Subsequently, we generated sequencing libraries for both the pancreatic islets and EndoC-βH1 samples using the Qiaseq miRNA library kit, followed by Illumina platform-based sequencing. The complete experimental process is delineated in **Fig. 1a**.

**Figure 1.**
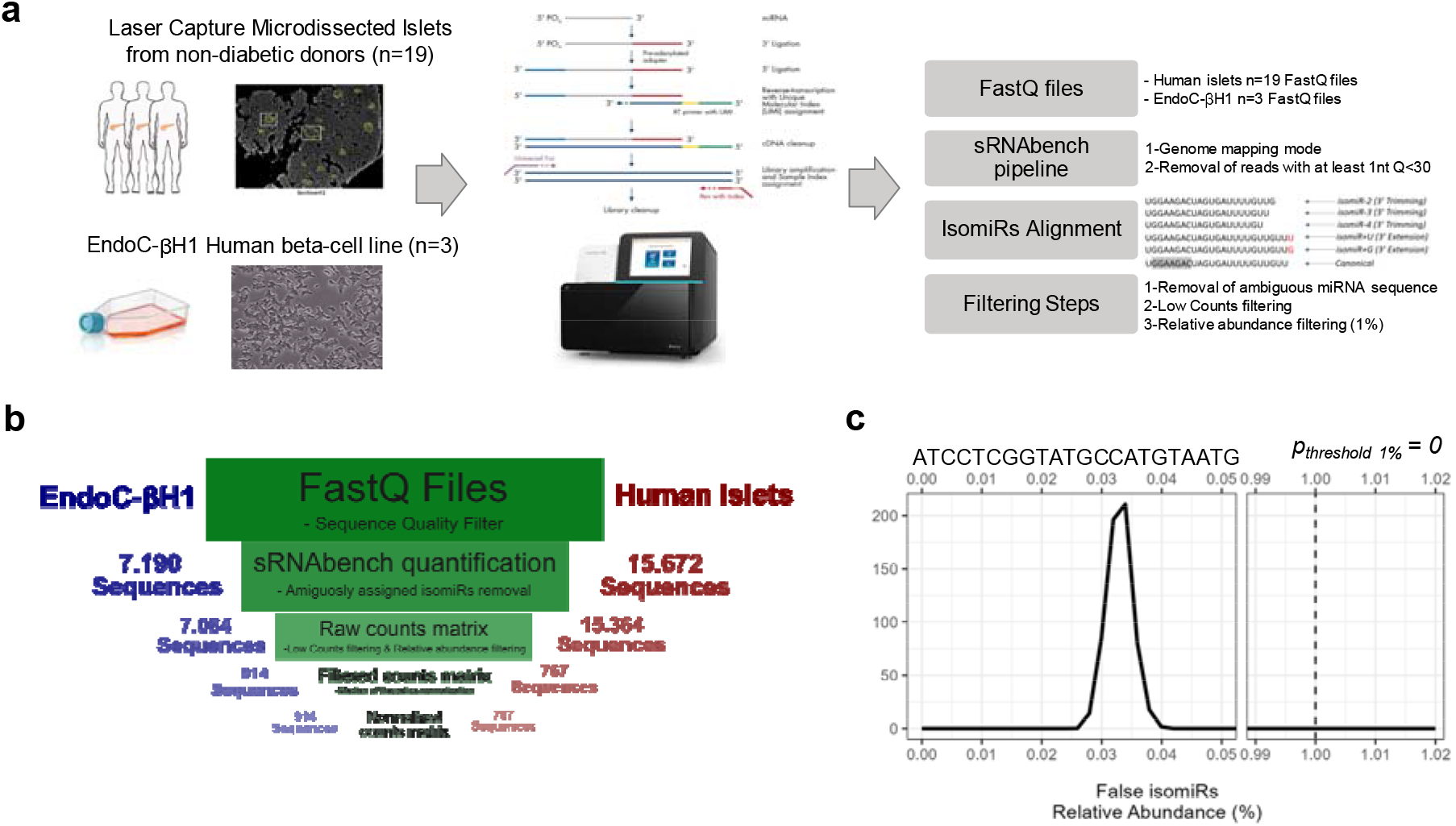
Study design and isomiR sequencing quality controls. **(a)** Study workflow scheme, from RNA extraction to isomiR bioinformatic pipeline. FastQ files were analyzed with the sRNAbench online pipeline in genome mapping mode and with the implementation of a sequence quality filter. After the alignment and the quantification step, additional filtering steps were implemented to remove ambiguous sequences, sequences with low expression (low-counts filtering) and sequences with a low contribution to total miRNA expression (relative abundance filtering). **(b)** Scheme showing the number of sequences detected in HI and EndoC-βH1 after (i) quantification, (ii) removal of ambiguously mapped squences, (iii) Low counts filtering & relative abundance filtering and (iv) Normalization. **(c)** Density plot from in-silico simulation shows the relative abundance distribution of false isomiRs sequences with a single nucleotide variation compared to the reference sequence (*ATCCTCGGTATGCCATGTAATG)*. The procedure was carried out simulating the reading the reference sequence 1.000.000 times, with a probability of error in the reading of each nucleotide = 0,1% (Q=30). The relative abundance threshold of 1% was validated by computing the probability of false isomiRs with a relative abundance above the cut-off (p=0).

Several small RNA library sequencing kits have demonstrated high accuracy in detecting isomiRs, with the Qiaseq miRNA library kit being one of the most reliable preparative library chemistry kits [9]. Nevertheless, to enhance the accuracy at single-nucleotide resolution, which is critical for generating insights into the biological functions of isomiRs, stringent filtering criteria during sequencing dataset generation are imperative. To this end, we established a hierarchical filtering process comprising, (*i*) an initial assessment of the read quality and (*ii*) the application of a relative abundance filter in conjunction with a low count filtering step. In LCM HI and EndoC-βH1 samples, an average of 31,1 ± 3,1% and 19,9 ± 1,2%, respectively, of adapter-trimmed reads were excluded via the initial quality filter. This preliminary filtering step eliminated a substantial portion of reads with at least one nucleotide possessing a Quality score below 30. Then, we removed ambiguously assigned isomiRs (sequences assigned to multiple miRNAs) and applied the relative abundance filter and the low counts filter (**Fig. 1b**). The effectiveness of the relative abundance filter in eliminating sequencing errors was assessed using an in-silico procedure involving the sequencing simulation of a 22-nucleotide sequence through the generation of 1L10^6^ reads, subjected to the typical experimentally observed single-nucleotide errors (Q=30). This evaluation indicated that the likelihood of false isomiRs with a relative abundance filter exceeding the threshold of 0.01 (i.e. 1 %) was estimated to be *p*=0, underscoring the filter’s efficacy in eliminating false isomiRs stemming from single nucleotide sequencing errors (**Fig. 1c**). Furthermore, this in-silico simulation validated the significance of the filtering step based on its contribution to the corresponding miRNA counts.

Finally, our bioinformatics pipeline allowed the identification of 767 and 914 unique, unambiguously assigned, and relevant miRNAs and their isoform sequences in HI and EndoC-βH1 samples, respectively.

### Human pancreatic islets and EndoC-βH1 miRNA-sequencing reveals a relevant fraction of 3’ isomiRs

MiRNA-sequencing analysis of LCM HI from n=19 non-diabetic subjects revealed a total of 767 sequences originating from 164 annotated miRNAs. Of these, n=31 miRNAs were present as canonicals and n=29 exclusively as isomiRs, while n=104 both as canonical and isomiRs (**Fig. 2a**). In EndoC-βH1 beta cells, we detected a total of 914 sequences originating from 243 annotated miRNAs. Of them, n=45 miRNAs were present as canonicals and n=50 exclusively as isomiRs, while n=148 both as canonicals and isomiRs (**Fig. 2b**).

**Figure 2.**
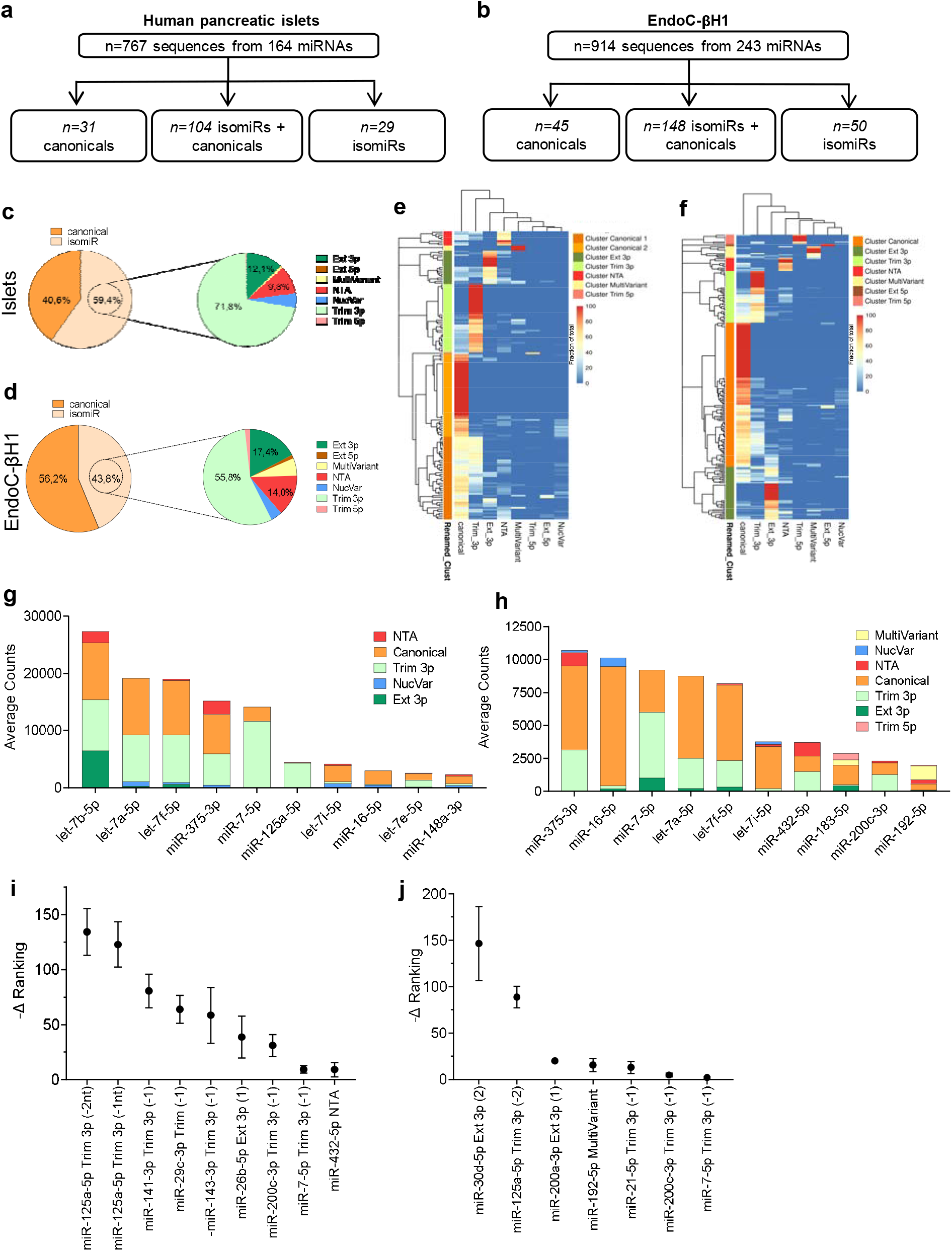
Expression Profiling of isomiRs in human pancreatic islets and EndoC-βH1. **(a,b)** Scheme showing the distribution of the sequences detected after the multiple filtering steps in human pancreatic islets (HI) (a) and in EndoC-βH1 (b). (**c,d**) Pie charts representing the average percentage of counts assigned to miRNAs and isomiRs and the detailed composition of the isomiR fraction in HI (c) and in EndoC-βH1 samples (d). (**e,f**) Heatmaps showing the results of the clustering analysis of the average composition of each miRNA across HI samples (e) and EndoC-βH1 (f); hierarchical clustering analyses were performed using the euclidean distance and the complete-linkage method. The optimal number of clusters was determined with the silhouette method, resulting in n=7 optimal clusters in HI. For each cluster identified, the name was determined based on the most prevalent isomiR class in its composition. (**g,h**) Barplots showing the isomiR types composition of the n=10 most expressed miRNAs in HI (g) and in EndoC-βH1 (h), computed as the average expression of each isomiR class across all samples. (**i,j**) Delta ranking analyses between specific isomiR sequences and related canonicals in HI (i) and EndoC-βH1 (j); in the scatterplots, the y-axis represents the -Delta Ranking between isomiR expression rank and the corresponding canonical miRNA expression rank. Only isomiR sequences, among the n=50 most expressed sequences, with an average rank lower than the corresponding canonical miRNA’s average rank are reported. High values of –Delta Ranking are related to isomiRs much more expressed than the corresponding canonical miRNA. For each reported isomiR, the mean and the standard deviation of the -Delta Ranking across samples are reported. Hi: n=19; EndoC-βH1: n=3 independent cell passages.

Both in HI and EndoC-βH1 cells, isomiR/canonical expression revealed a surprisingly high proportion of counts assigned to isomiRs in comparison to canonical miRNA sequences, detecting 59,4% and 43,8% of isomiR counts accounting to total miRNAs expression respectively in HI and EndoC-βH1 cells (**Fig. 2c** and **2d**). A more in-depth analysis of isomiRs expression was based on the different sequence variation types, or classes. In particular, according to the modification, sequences were defined as 5’ or 3’ end trimming (Trim 5p and Trim 3p), 5’ or 3’ end extension (Ext 5p and Ext 3p), Non-Templated Addition (NTA), multi-length variant (MultiVariant), Nucleotide Variation (NucVar) and canonical (see ESM Methods). The analysis revealed that the most prevalent class of isomiRs is represented by Trim 3p (HI: 71,8%; EndoC-βH1: 55,8%), followed by Ext 3p (HI:12,1%; EndoC-βH1:17,4%), NTA (HI: 9,8%; EndoC-βH1: 14,0%), NucVar (HI: 5,1 %; EndoC-βH1: 4,0 %), MultiVariant (HI: 0,7 %; EndoC-βH1 5,9%) and by a minor fraction of 5’ end variations in both HI and EndoC-βH1 (**Fig. 2c** and **2d** and **Table 1**). To eliminate the possibility that even a portion of Trim 3p isomiRs might originate from random RNA degradation, we conducted a correlation analysis between the quantity of Trim 3p and the RNA Integrity Number (RIN) measured across all LCM-HI samples. The result revealed no significant correltion, thus excluding any association between Trim 3p levels and RNA degradation (**ESM Fig. 1**).

While some miRNAs in both HI and EndoC-βH1 cells are exclusively present as canonical or isomiR forms (**Fig. 2a** and **2b**), many of them exhibit a unique pattern characterized by a heterogeneous sequences composition. This heterogeneity enabled us to categorize miRNAs based on their composition pattern in terms of canonical/isomiRs categories, thus facilitating the identification of distinct clusters of miRNAs that share similar patterns and most likely subjected to the same post-transcriptional regulatory mechanisms (**Fig. 2e** and **2f**). Hence, in HI, the hierarchical clustering analysis revealed n=7 distinct clusters of miRNAs: Cluster Canonical 1 (n=47 miRNAs; 51.3% of canonicals followed by 28.5% Trim 3p), Cluster Canonical 2 (n=47 miRNAs; 91.8% of canonicals), Cluster Ext 3p (n=19 miRNAs; 70.5% of Ext 3p and 12.5% of canonicals), Cluster MultiVariant (n=3 miRNAs; 100% of MultiVariant), Cluster NTA (n=8 miRNAs; 63.8% of NTA followed by 24.3% of canonicals), Cluster Trim 3p (n=39 miRNAs; 85.6% of Trim 3p and 9.2% of canonicals) and Cluster Trim 5p (n=1 miRNA; 68.6% of Trim 5p and 31.4% of canonicals) (**Fig. 2e**).

In EndoC-βH1 we observed a similar miRNA patterning with the identification of n=7 clusters. Similarly to HI, the most numerous cluster was the “Canonical” (n=123 miRNAs; 79.4% of canonicals, 10.0% of Trim 3p), followed by Cluster Ext 3p (n=45 miRNAs; 67.4% of Ext 3p and 14.1% of canonicals), Cluster Ext 5p (n=2 miRNA; 100% Ext 5p), Cluster MultiVariant (n=10 miRNAs; 73.0% of MultiVariant and 9.6% of canonicals), Cluster NTA (n=11 miRNAs; 79.7% of NTA followed by 12.9% of canonicals), Cluster Trim 3p (n=44 miRNAs; 71.2% of Trim 3p and 22.1% of canonicals) and Cluster Trim 5p (n=8 miRNA; 86.5% of Trim 5p and 7.6% of Trim 3p) (**Fig. 2f**). It is noteworthy that the internal composition of each cluster exhibited a significant level of concordance between HI and beta cells (**ESM Fig. 2**). As expected, canonical clusters showed the highest concordance rate between HI and EndoC-βH1 (n=68 miRNAs shared and a Jaccard similarity index of 0.78) (**ESM Fig. 2** and **ESM Table 4**). Within the isomiR clusters, Trim 3p clusters shared 26 miRNAs with a Jaccard similarity index of 0.78, thus exhibiting a noteworthy concordance between HI and EndoC-βH1 cells (**ESM Fig. 2** and **ESM Table 4**). Among isomiRs included in this cluster, we identified some miRNAs that are highly expressed in HI and beta cells and have previously shown important functions (i.e. miR-200 family[42], miR-7 [43, 51], miR-29 [52], miR-21 [53]). This suggests that some crucial miRNAs in beta cell or in HI predominantly exist as isoforms rather than canonical sequences.

**Table 4.**
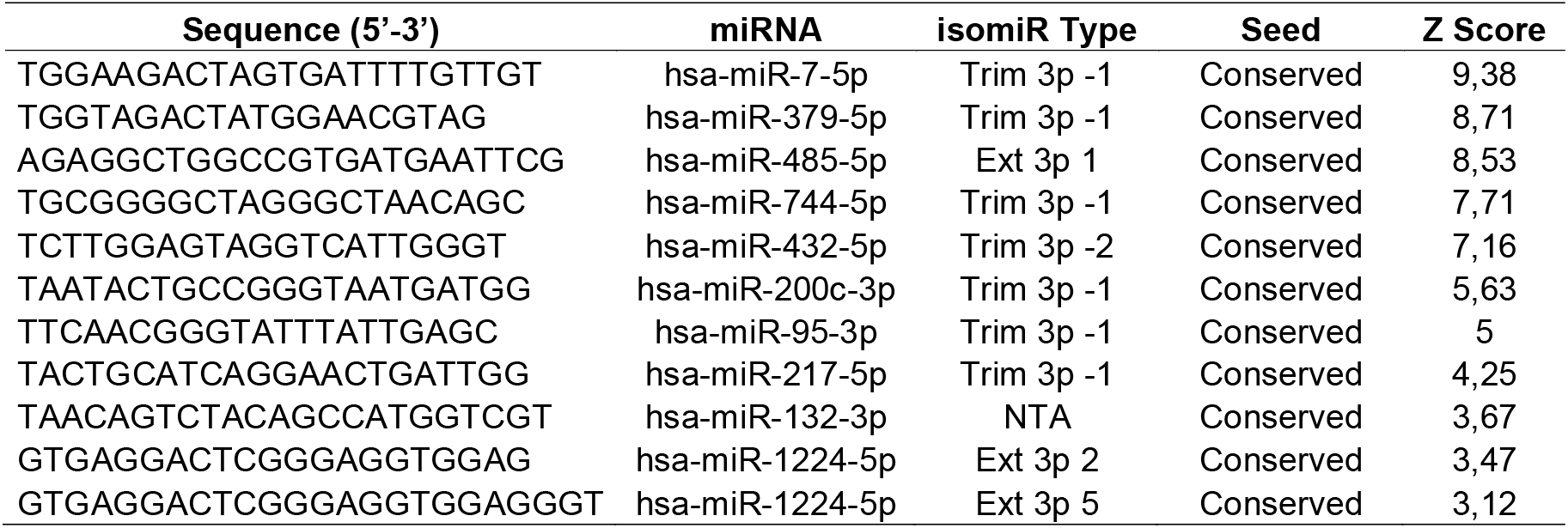
Beta cell enriched isomiRs. Main characteristics of the beta cell enriched isomiRs according to isomiRdb analysis (Z score>3). Sequence, miRNA of origin and isomiR type (with the number of nucleotides trimmed or added) are shown. Conservation of the isomiR seed sequence respect to the canonical miRNA is also reported (conserved/not conserved). Z score showed the enrichment in beta cells respect to the other cell type (n=99) included in isomiRdb. IsomiRs are shown from the most enriched to the least, among those showing a Z-score >3.

These findings prompted us to delve deeper into the specific sequence composition of the most highly expressed miRNAs in HI and EndoC-βH1 cells. Notably, when we analyzed the composition of the n=10 most expressed miRNAs, we observed a significant contribution of isomiRs to the overall expression of these highly represented miRNAs. Additionally, there was a notable heterogeneity in the abundance of isomiR variants, which appeared to be miRNA-dependent (**Fig. 2g** and **2h**). For instance, in HI, the expression of let-7b-5p, which was the most highly expressed miRNA, stemmed from 64% of isomiR variants and 36% of canonical sequence (**Fig. 2g** and **Table 2**). In EndoC-βH1 cells, miR-375-3p, the most highly expressed miRNA, was composed by a significant fraction of isomiRs (40% isomiRs and 60% canonical) (**Fig. 2h**) and a similar pattern was observed in HI (55% isomiRs and 45% canonical). Strikingly, we observed that the expression of miR-7-5p, a miRNA enriched in beta cells [43], was predominantly composed of isomiR sequences in both HI (82%) and EndoC-βH1 (65%), rather than canonical ones (**Fig. 2g, 2h**, and **Table 2, 3**); of note, the isomiRs of miR-7-5p were primarily characterized by Trim 3p, with some sequences exhibiting significant truncation of the 3’ end (−4 nt), a pattern observed in both HI and EndoC-βH1 cells (**ESM Fig. 3**).

**Figure 3.**
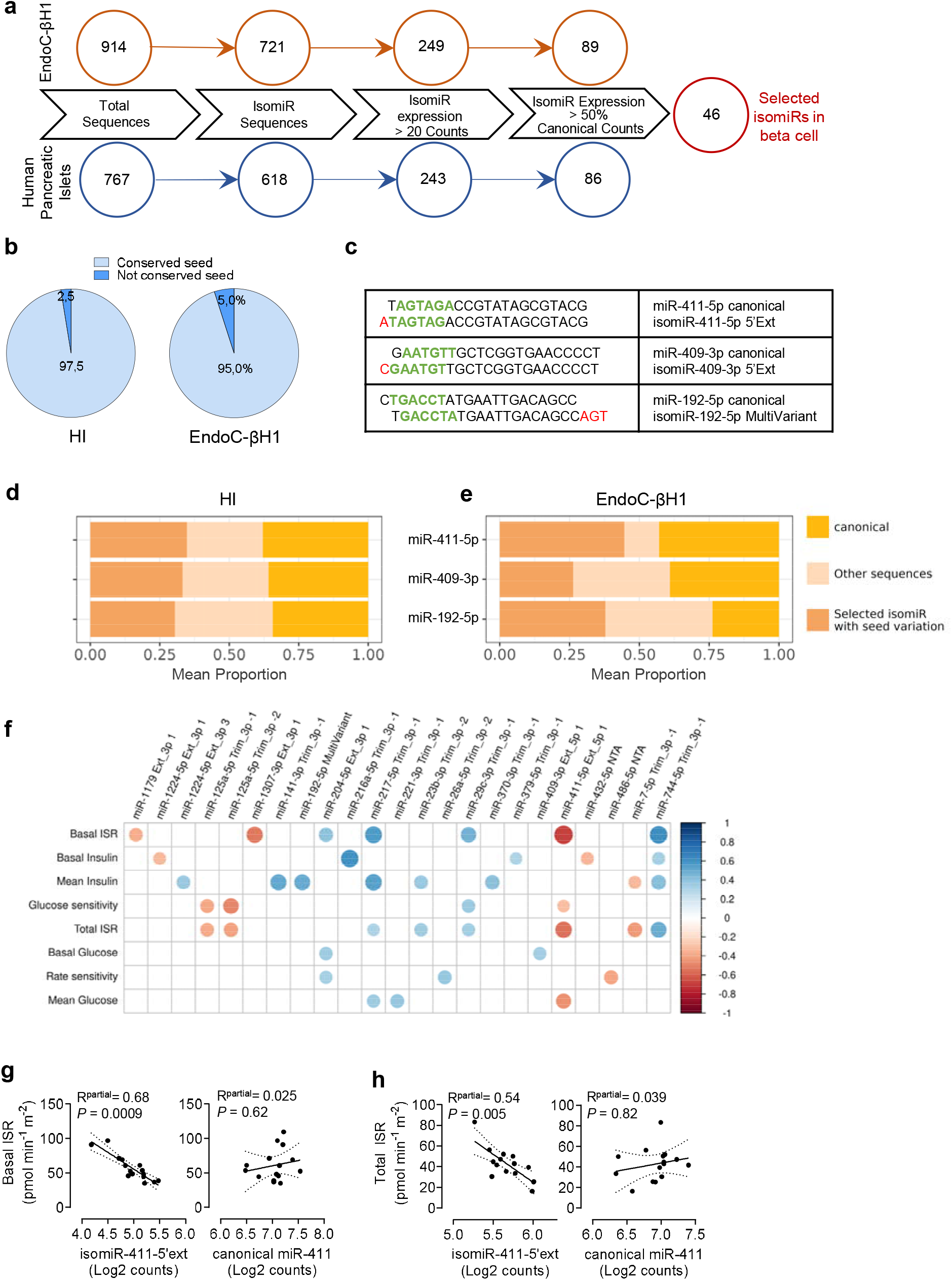
Identification of a beta cell isomiRs signature. (**a**) Scheme reporting the filtering steps for the identification of a signature of isomiRs robustly detected in both HI and EndoC-βH1. The number of sequences after each filtering step in EndoC-βH1 and HI are reported in the graph. (**b**) Pie charts showing the average contribution of isomiRs with seed variation to total miRNA expression across EndoC-βH1 and HI samples. (**c**) Detailed sequences of the 3 isomiR (isomiR-411-5p 5’Ext, isomiR-409-3p 5’Ext, isomiR-192-5p MultiVariant) with seed changes and their canonical counterparts. The seed sequence is represented in green, while nucleotide modifications are reported in red. (**d,e**) Barplots showing the average expression composition of the miR-411-5p, miR-409-3p and miR-192-5p in HI (d) and in EndoCβH1 (e); the average contribution of the selected isomiR sequence, the canonical sequence and all the other isomiR sequences are reported in different colors. (**f**) Correlation plot showing significant associations between signature isomiRs and OGTT-derived metabolic parameters in NGT living individuals. Data were corrected for age, gender and BMI. The square root of the partial *R2* of the multiple linear regression is shown to visualize positive (blue) or negative (red) significant associations (*P*≤0.05). Circle size is correlated with the magnitude of the p-value. Only significant results in at least one isomiR and a clinical parameter are reported. (**g,h**) Scatterplot showing the association of isomiR-411 5’ Ext and its canonical counterpart expression with (g) basal insulin secretion rate (Basal ISR) and with (h) total insulin secretion rate (Total ISR) measurements available from n=15 living donors. IsomiR and canonical miRNA expression are reported as normalized reads counts (log2 scale), corrected for the effect of the covariates (age, gender and sex). The p-value of the clinical parameter’s coefficient obtained in multiple linear regression model and the square root of its partial R^2^ are reported.

To confirm the existence of isomiR sequences that are more highly expressed than their respective canonical counterparts, we conducted a ranking analysis among the n=50 most expressed sequences. In HI, we detected n=9 isomiRs showing a higher expression respect to their canonical counterparts (**Fig. 2i**), while n=7 isomiRs were detected in EndoC-βH1 cells (**Fig. 2j**). In HI, we noticed that the most expressed isomiR in comparison to its canonical counterpart, was miR-125a-5p Trim 3p (both 2nt and 1nt) (**Fig. 2i**). Additionally, we found that the 3’ trimming isomiR of miR-125a-5p was also present in EndoC-βH1 cells, where it ranked as the second most expressed isomiR relative to its canonical counterpart (**Fig. 2j**).

Collectively, these results demonstrate the significant contribution of isomiRs to the global expression profiles of miRNAs in HI and EndoC-βH1 cells.

### A set of isomiRs showed variations in canonical seed sequences with potential implications for beta cell function

To identify a robust set of isomiRs consistently expressed both in HI and EndoC-βH1 cells and representing a relevant fraction of the total miRNA counts, we conducted a comprehensive datasets cross-validation and intersection. We reasoned that if a particular isomiR sequence is detected in both HI and EndoC-βH1 at noteworthy level, it is likely expressed in beta cells where it may potentially hold a functional role. To achieve this, we implemented consecutive filtering steps to select highly expressed isomiRs sequences with an average representation of ≥50% of corresponding canonical miRNA counts and an average expression >20 counts (**Fig. 3a** and ESM Methods). We identified n=46 isomiRs expressed both in HI and EndoC-βH1 and showing the above reported characteristics (**Fig. 3a** and **ESM Table 5**).

Since the functional significance of isomiRs respect to their canonical counterparts is primarily attributed to alterations in the seed sequences, our focus was on those isomiRs displaying such changes, particularly from Trim 5p, Ext 5p, NucVar, or MultiVariant isomiRs. The analysis of the complete datasets showed that a minor fraction of isomiR sequences exhibited seed changes, accounting for 2,5% in HI and 5,0% in EndoC-βH1 cells (**Fig. 3b**). Focusing on the set of previously selected 46 beta cell isomiRs (**ESM Table 5**), we identified three of them exhibiting seed sequence changes compared to their canonical counterparts: isomiR-411-5p Ext 5p, isomiR-409-3p Ext 5p and isomiR-192-5p MultiVariant (**Fig. 3c**). These sequences were notable for accounting to a significant proportion of the overall miRNA expression both in HI and EndoC-βH1 (isomiR-411-5p Ext 5p: 35% HI, 45% EndoC-βH1; isomiR-409-3p Ext 5p: 33% HI, 26% EndoC-βH1; isomiR-192-5p MultiVariant: 30% HI, 38% EndoC-βH1) (**Fig. 3d** and **3e**). This observation strongly suggests that they may have distinct functions compared to their canonical counterparts.

Finally, in order to explore the potential function of these isomiRs in beta cells, we conducted a correlation analysis between the isomiRs signature and a set of parameters obtained from in vivo metabolically profiled NGT living donors (**ESM methods**). Remarkably, among the isomiRs displaying seed sequence changes, isomiR-411-5p Ext 5p, but not its canonical counterpart, exhibited a major significant inverse correlation with beta cell functional parameters (**Fig. 3f**), including basal insulin secretion rate and total insulin secretion rate (**Fig. 3g** and **3h**). These results highlight the potential alternative functional role of isomiRs compared to their canonical counterparts and underline the importance to evaluate them and their related functions.

### Validation of the beta cell isomiRs signature

To validate the identification and expression pattern of this set of n=46 isomiRs signature, and to investigate their potential specificity or enrichment within human beta cells, we conducted a comprehensive analysis of an external database that provide access to consistently analysed isomiRs from small RNA sequencing data obtained from n=187 distinct cell types and n=52 different human tissues (isomiRdb), including sorted human primary beta cells [54]. As isomiRdb contains data from both healthy and diseased tissues and cell types, we firstly filtered out experiments related to unhealthy conditions, to focus exclusively on datasets with physiological isomiR expression profiles. This resulted in a total of n=99 different human cell types. Subsequently, we implemented a multi-step filtering process to obtain a reliable set of isomiRs from these datasets. Finally, we verified the detection, expression, and beta cell specificity of our n=46 isomiR signature (**Fig. 4a**) in the curated isomiRdb dataset.

**Figure 4.**
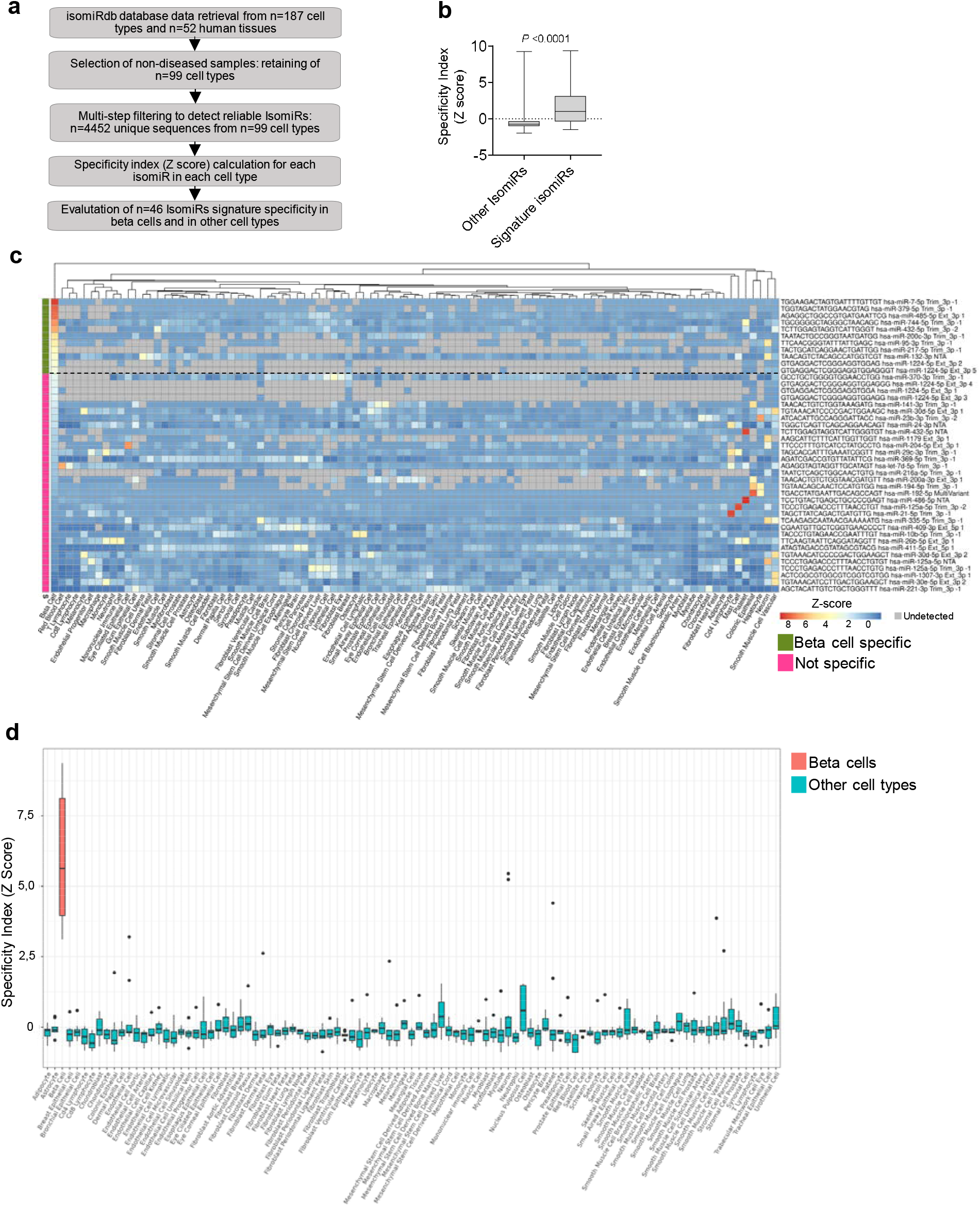
Validation of the beta cell isomiR signature. (**a**) Scheme reporting the pipeline adopted for the analysis of the beta cell isomiR signature in isomiRdb database. (**b**) Boxplot showing the specificity index (Z-score) of the n=43 beta cell signature’s isomiRs analysed in isomiRdb, compared to the specificity index from the other isomiRs; Wilcoxon test was performed to evaluate the significance of the differences in Z-scores, considering as significant *P* ≤ 0.05. (**c**) Heatmap reporting the specificity indexes (Z-scores) of the 43 beta cell signature isomiRs sorted in a descending order respect to their enrichment in beta cell (from the most to the least enriched). A hierarchical clustering analysis for cell populations was performed (euclidean distance and complete agglomeration method). The dashed line represents the threshold cutoff corresponding to Z-score=3, used for the identification of the beta cell enriched isomiRs (n=11). Z-score values are reported as scale color from blue (low specificity) to red (high specificity). Grey boxes indicate *not detected* isomiR expression. (**d**) Boxplot of the n=11 isomiRs from the signature detected as specific (Z-score>3) among the different cell types present in isomiRdb. Beta cell are shown in red, while all the other cell types in turquoise.

Out of the previously identified n=46 beta cell isomiRs, n=43 were detected in isomiRdb datasets and expressed in at least one cell type, thereby validating the authenticity of the identified sequences. To assess the enrichment of these n=43 isomiRs in beta cells compared to other isomiRs, we computed a specificity score (Z-score) to measure their unique expression patterns in each cell type. Notably, within beta cells, our signature exhibited a significantly higher specificity index (mean Z-Score=1.9) compared to other isomiRs (mean Z-score= −0.48) (*P*<0.0001) (**Fig. 4b**). However, we observed that not all n=43 isomiRs contributed equally to the overall specificity of the signature. Specifically, n=11 isomiRs (**Table 4**) retained a Z-score >3, indicating their particularly high specificity in beta cells. This was confirmed by heatmap clustering analysis, which showed the elevated enrichment of these n=11 isomiRs in beta cells in comparison to all other cell types (**Fig. 4c**). Furthermore, when we considered the specificity indexes from this set of isomiRs, it further corroborates the high specificity observed in beta cells compared to all other cell types (**Fig. 4d**).

Collectively, these data demonstrate two key findings: (*i*) the authenticity of n=43 isomiR sequences detected in our datasets, including isomiR-411-5p, −409-3p and −192-5p, being confirmed in independent external sequencing experiments, and (*ii*) the notably high specificity of a set of n=11 isomiRs within beta cells compared to other cell types.

## Discussion

IsomiRs are considered non-artifactual variants of miRNA sequences [9] with specific functional roles in different contexts. Notably, 5’ isomiRs feature a shifted seed sequence compared to canonical miRNAs, leading to significant alterations in their targetome. Similarly, although less well-defined, 3’ isomiRs may have distinct non-canonical target recognition and, importantly, exhibit differences in half-life and AGO2 loading. In both instances, isomiRs exhibit alternative functions compared to their canonical counterparts.

Here, in light of their functional role, we conducted a comprehensive analysis of isomiRs in human pancreatic islets (HI) from non-diabetic living donors and the human beta cell line EndoC-βH1. Our goal was to provide a detailed overview of isomiRs expression in beta cells and to explore potential functional implications in this context. To achieve this, we utilized a small RNA sequencing approach, employing an up-to-date and reliable miRNA-seq library protocol. We combined this with a state-of-the-art bioinformatic pipeline for isomiR detection (sRNAbench) which was implemented with stringent multi-step filtering. Additionally, the analysis of laser capture microdissected islets from living donors fully metabolically characterized allowed us to unprecedently associate isomiRs expression with beta cell in vivo functional parameters and glucose homeostasis measurements.

Firstly, our findings revealed that a significant proportion of total miRNA counts, both in HI and EndoC-βH1, consisted of 3’ end isomiRs with trimming modifications as a prevalent component. Of note, 3’end isomiRs expression was not related to the overall RNA integrity variations in LCM HI samples. Importantly, we observed a high concordance rate in terms of specific miRNAs showing a prevalent Trim 3p fraction in both HI and EndoC-βH1, with 26 miRNAs displaying a highly similar composition pattern. This suggests that the generation of 3’ end isomiRs through trimming may be highly regulated and sequence-specific. Indeed, we found that in HI, miR-7a-5p was primarily composed of Trim 3p isomiRs, with some sequences trimmed by up to 4 nucleotides, and this composition profile was also observed in EndoC-βH1, confirming the presence of a conserved mechanism of post-transcriptional miRNA regulation. Interestingly, this unique pattern was not observed in other miRNAs, even those with higher expression levels than miR-7 (e.g., miR-375-3p or let-7b-5p). This finding demonstrates that these post-transcriptional modifications are specific to individual miRNAs and are not correlated with their expression levels.

Secondly, we demonstrated that some specific isomiR sequences are more expressed that the canonical one which was commonly identified as the main (and unique) represented feature in most cases and studies. Interestingly, among others, we identified miR-125a-5p Trim 3p isomiR as more expressed respect to its canonical both in HI and EndoC-βH1. This finding implies putative functional differences that should be addressed in future studies in light of the high expression of this isomiR.

Thirdly, by intersecting and cross-validating HI and EndoC-βH1 dataset, we obtained a set of n=46 reliably expressed isomiRs. Among them, we identified three isomiRs with seed sequence variation (miR-411-5p, miR-409-3p, miR-192-5p), having relevant functional implications based on the changes in their targetome. Indeed, a correlation analysis between isomiRs expression in HI and in-vivo metabolic clinical parameters measured on the same non-diabetic subjects, showed significant correlations between their expression levels and specific parameters of beta cell functions. The most relevant resulted isomiR-411-5p Ext 5p which is negatively correlated with both basal insulin secretion rate and total insulin secretion rate, while the same correlation was not observed for its canonical sequence, thus demonstrating the importance to evaluate isomiRs in the context of beta cell function.

Finally, we validated the set of 46 isomiR sequences in external datasets included in the isomiRdb repository and composed of small RNA sequencing experiments from multiple sources. Such analysis allowed us to confirm the bona-fide of the isomiR sequences identified, and to define a set of n=11 isomiRs significantly associated to beta cells and which showed a higher specificity score respect to other human cell types.

The molecular mechanisms responsible for the generation of isomiRs in beta cells warrant further investigation. Numerous reports have suggested that 3’ isomiRs, the most prevalent class in beta cells, may be generated through the action of 3’-5’ exoribonucleases, such as Nibbler (Nbr). Notably, in *Drosophila Melanogaster*, the specific knockout of the Nbr exoribonuclease leads to the loss of Trim 3p isomiRs, resulting in critical dysfunctions in flies and underscoring their functional significance [55][56]; Interestingly, Nibbler homologs include EXD3 gene in human, thus suggesting a similar regulation in human cells as well. Another enzyme involved in exoribonuclease activity of miRNA is PARN which should be taken into consideration, alongside EXD3, as a potential factor in the generation of 3’IsomiRs in beta cells [57].

In contrast, the biogenesis of 5’ isomiRs in the beta cell context appears to be more selective and finely regulated than that of 3’ isomiRs, as they are represented by a smaller number of miRNAs. Notably, 5’ isomiRs exhibit changes in the miRNA seed region, resulting in a different set of mRNA targets respect to the canonical. This likely leads to an alternative functionality, as supported by our findings, which demonstrate a strong correlation between isomiR-411-5p Ext 5p, but not its canonical, and basal beta cell functions. The regulation of isomiR-411 generation and whether it is influenced by metabolic or inflammatory stresses should be further elucidated.

In conclusion, we have reported an overview of the isomiR profile in LCM-human pancreatic islets and Endoc-βH1 cells, thus establishing the first comprehensive set of isomiRs associated with beta cells. Our study provides a solid foundation for conducting further analyses to investigate the functional roles of isomiRs in both normal and diabetic conditions.

## Supporting information

ESM file

## ACKNOWLEDGEMENTS

The secretarial help of Alessandra Mechini and Maddalena Prencipe was highly appreciated.

## DATA AVAILABILITY STATEMENT

Raw and analysed data are available from the corresponding author upon request.

## CONTRIBUTION STATEMENT

SA and EA conducted the experiments and the analyses, prepared tables and figures, interpreted the data and co-wrote the manuscript. MT, GEG, DF, GL, MB, AM, AB evaluated, discussed and interpreted the data. AG and TM coordinated the access to pancreatic tissues and the full in vivo metabolic profiling and interpreted the data. GP and VT performed pancreatic surgery and collected pancreatic samples. GDG and LS performed living donors recruitment and metabolic evaluation, AM performed mathematical modelling analysis of OGTT data, FD and GS initiated, designed and supervised the study, interpreted the data and wrote and revised the manuscript. SA, EA, FD and GS are the guarantors of this work and, as such, had full access to all the data in the study and takes responsibility for the integrity of the data and the accuracy of the data analysis.

## Abbreviations

HI: Human Islets
LCM: Laser Capture Microdissection
NTA: Non-templated addition
NucVar: Nucleotide Variation
MultiVariant: Multi-length Variant Trim
3p/5p: 3’/5’ end Trimming
Ext 3p/5p: 3’/5’ end Extension

