## Supplementary material for "Comprehensive IsomiR sequencing profile of human pancreatic islets and EndoC-βH1 beta-cells": ESM file

**Electronic Supplementary Material (ESM)**

**ESM METHODS**

**Study subjects and human pancreas tissue collection**

Human pancreatic sections analyzed in this study were obtained from pancreata of n=3 brain-dead adult non-diabetic multiorgan donors within the European Network for Pancreatic Organ Donors with Diabetes (EUnPOD) and from n=16 non-diabetic living donors undergoing pylorus-preserving pancreatoduodenectomy recruited at the Digestive Surgery Unit and studied at the Centre for Endocrine and Metabolic Diseases Unit (Agostino Gemelli University Hospital, Rome, Italy). Indications for surgery were periampullary tumors, pancreatic intraductal papillary tumors, mucinous cystic neoplasm of the pancreas, and nonfunctional pancreatic neuroendocrine tumors. (**ESM Table 1**). Surgical pancreatic tissue specimens were snap frozen in liquid nitrogen and stored at -80°C embedded in Tissue-Tek OCT compound.

The study protocol (ClinicalTrials.gov NCT02175459) was approved by the Ethical Committee Fondazione Policlinico Universitario Agostino Gemelli IRCCS – Università Cattolica del Sacro Cuore (P/656/CE2010 and 22573/14), and all participants provided written informed consent, followed by a comprehensive medical evaluation, as previously described [[1, 2]](https://sciwheel.com/work/citation?ids=15612984,13977332&pre=&pre=&suf=&suf=&sa=0,0&dbf=0&dbf=0).

In INNODIA EUnPOD network, pancreata not suitable for organ transplantation were obtained with informed written consent by organ donors’ next-of -kin and processed with the approval of the local ethics committee of the Pisa University (**ESM** **Table 2**).

None of the patients enrolled had a family history of diabetes. Living donors patients underwent both a 75-g oral glucose tolerance test and glycated hemoglobin (HbA1c) testing to exclude diabetes, according to the American Diabetes Association criteria [[3]](https://sciwheel.com/work/citation?ids=12466885&pre=&suf=&sa=0&dbf=0).

**Oral Glucose Tolerance Test**

A standard 75 g oral glucose tolerance test was performed after a 12h overnight fast with measurements of glucose, insulin and C-peptide at 0, 30, 60, 90, 120 min after the glucose load.

Based on OGTT, we classified the population as normal glucose tolerant (NGT) if 2-hour post load glucose was below 140 mg/dl [[3]](https://sciwheel.com/work/citation?ids=12466885&pre=&suf=&sa=0&dbf=0).

During OGTT, insulin secretion was derived from C-peptide levels by deconvolution. Betacell glucose sensitivity (GS), i.e., the slope of the relationship between insulin secretion and glucose concentration, was estimated from the OGTT by modelling, as previously described [[4, 5]](https://sciwheel.com/work/citation?ids=11524219,9771328&pre=&pre=&suf=&suf=&sa=0,0&dbf=0&dbf=0). Rate sensitivity (RS), also estimated from OGTT modelling, is a beta cell functional parameter that represents the dependence of the insulin secretion rate (ISR) on the rate of change in glucose concentration and is related to early insulin release.

**Laser Capture Microdissection**

Pancreatic human tissue samples n=19 from non-diabetic donors (**ESM** **Table 1** and **ESM Table 2**) were frozen in Tissue-Tek OCT compound and then 7-μm thick sections were cut from frozen O.C.T. blocks. Sections were fixed in 70% Ethanol for 30 s, dehydrated in 100% Ethanol for 1 min, in 100% Ethanol for 1 min, in Xylene for 5 min and finally air-dried for 5 min. Laser Capture Microdissection (LCM) was performed using an Arcturus XT Laser-Capture Microdissection system (Arcturus Engineering, Mountain View, CA, USA) by melting thermoplastic films mounted on transparent LCM caps (cat. LCM0214 - ThermoFisher Scientific, Waltham, MA, USA) on specific islet areas. Intrinsic beta cells autofluorescence allowed the identification of human pancreatic islets for LCM procedure. Adhesive thermoplastic caps containing microdissected cells were incubated with 10 μl of Extraction Buffer (cat. kit0204 - ThermoFisher Scientific, Waltham, MA, USA) for 30 min at 42 ◦C and kept at −80°C until RNA extraction. Each microdissection was performed within 30 min from the staining procedure in a contamination-free-dehumidified environment with an external temperature of 16°C to preserve RNA integrity. Overall, an endocrine mass of approximately 2x10^6^ µm^2^ area for each case was collected for further molecular analysis.

**EndoC-βH1 cell culture**

EndoC-βH1 human beta-cell line [[6]](https://sciwheel.com/work/citation?ids=1914391&pre=&suf=&sa=0&dbf=0) was obtained by UniverCell-Biosolutions (Toulouse-France) and used for all experiments between passages 78-88. EndoC-betaH1 were cultured at 37 °C with 5% CO2 in coated flask (coating medium: DMEM high-glucose cat. 51441C, Penicillin/Streptomycin 1% cat. P0781, ECM 1% cat. E1270 and Fibronectin from bovine plasma 0.2% cat. F1141 - all from Sigma Aldrich, St. Louis, MO, USA) and maintained in culture in low-glucose DMEM (cat. D6046) supplemented with 2% BSA fraction V (cat. 10775835001), β- Mercaptoethanol 50 μM (cat. M7522), L-Glutamine 1% (cat. G7513), Penicillin/Streptomycin 1%

(cat. P0781), Nicotinamide 10 mM (cat. N0636), Transferrin 5.5 μg/mL (cat. T8158) and Sodium selenite 6.7 ng/mL (cat. S5261) (all from Sigma Aldrich, St. Louis, MO, USA).

**RNA isolation and quality control**

Total RNA was extracted from each LCM sample using PicoPure RNA isolation kit Arcturus (cat. kit0204 - ThermoFisher Scientific, Waltham, MA, USA) following manufacturer’s procedure. Briefly, the cellular extracts were firstly mixed with 12.5 μl of 100% Ethanol to obtain an enrichment also in small RNAs and then transferred onto the purification column filter membrane. DNase treatment was performed using RNase-Free DNase Set (cat. 79254 - Qiagen, Hilden, Germany). Total RNA was finally eluted in 11 μl of Elution Buffer (DNase/RNase-Free Water). All LCM captures deriving from the same human sample were pooled and subjected to a subsequent concentration through Savant SpeedVac™ DNA130 centrifugal evaporator. Total RNA was extracted from approximately 3.0 × 105 EndoC-βH1 through Direct-zol RNA Miniprep Kit (cat. R202-Zymo Research, Irvine, CA, US) following manufacturer’s instructions. Briefly, the pelleted cells were resuspended in QIAzol (cat. 79306, Qiagen), mixed with equal volume of 100% Ethanol and transferred to Zymo-Spin™ IICR Column. DNase digestion was performed using RNase-Free DNase Set (cat. 79254). Finally, RNA was eluted in 30 µl of DNase/RNase-Free Water.

In order to evaluate RNA abundance and purity, Agilent 2100 Bioanalyzer technology with RNA Pico chips (cat. 5067-1513 Agilent Technologies, Santa Clara, CA, USA) was performed for each RNA sample reporting RNA integrity (RIN) and concentration and by excluding samples with RIN<5.0 (RIN and concentration values for each sample are shown in **ESM Table 3**).

**Small RNA sequencing**

Small RNA seq was performed using QiaSeq miRNA library kit coupled with Illumina platforms sequencing as previously reported [[7]](https://sciwheel.com/work/citation?ids=11247261&pre=&suf=&sa=0&dbf=0). A total of 1 ng of RNA from microdissected human pancreatic Islets and from EndoC-βH1 was used to generate cDNA libraries using QiaSeq miRNA library kit (Qiagen, Hilden, Germany) following manufacturer’s instructions. QiaSeq library kit chemistry assigns Unique Molecular Index (UMI) during reverse transcription step to every small RNA molecule, to enable unbiased and accurate small RNAome-wide quantification. cDNA from Small RNA libraries were amplified and purified using magnetic beads. Then, libraries quality control (QC) was performed measuring cDNA concentration through QUBIT 3.0 spectrofluorometer (Qubit™ dsDNA HS Assay Kit, ThermoFisher Scientific, Waltham, MA, USA) and assessing their quality using capillary electrophoresis in Bioanalyzer 2100 (Agilent High Sensitivity DNA kit cat. 5067-4626). High quality of libraries was evaluated considering electropherograms showing a peak comprised between 175 and 185 bp.

Following libraries QC, all were normalized to 2 nM and pooled, denatured in 0.2 N NaOH and sequenced on Illumina NovaSeq 6000 platform (NovaSeq 6000 SP Reagent Kit v1.5 (100 cycles) using the XP protocol applying 75x1 single reads, or Illumina NextSeq550 platform (NextSeq 500/550 Mid Output Kit v2.5 (150 cycles), applying 75x1 single reads).

**sRNAbench pipeline for small RNA-seq**

FastQ files obtained from the sequencing were analyzed with the sRNAbench online pipeline [[8]](https://sciwheel.com/work/citation?ids=15387713&pre=&suf=&sa=0&dbf=0). Reads were processed with the Qiagen (with UMIs) protocol for adapters and duplicates removal.

Being interested in isomiRs analysis, a strong quality control step was implemented to ensure a high quality of each base in the reads. Using the minimum quality score threshold per sequence nucleotide, all reads with at least 1 nucleotide with a quality score (Q) < 30 (Phread+33) were discarded from the analyses. Reads were mapped in genome mapping mode, using the Human reference Genome Reference Consortium Human Build 38 patch release 13 (GRCh38.p13). Alignment was performed with the bowtie algorithm using the seed option with length (L) = 20, maximum number of mismatches (N) = 1, minimum length of the read =15, minimum number of read count =2 and maximum number of multiple mappings equal to 10. The seed alignment options of bowtie allow a maximum of N (1) mismatches in the first L (20) nucleotides, thus accounting for the great variability affecting the 3’ end of miRNAs. Mapped reads were annotated to miRNAs (and isomiRs) using miRBase release 22.1.

**IsomiR analysis and filtering steps**

The raw isomiRs counts matrix was obtained from the *microRNAannotation* file of each sample generated from sRNAbench. Sequences assigned to multiple miRNAs (usually resulting from Trim 3p) were removed from the counts matrix. However, some canonical miRNAs sequences can be assigned to multiple miRNAs (i.e., miR-199a and miR-199b, having the same sequence). To avoid the removal of these sequences, only ambiguous isomiRs (isomiRs assigned to multiple miRNAs) whose canonical counterpart was assigned to a single miRNA were removed. In this way, canonical miRNAs with the same sequence (i.e., miR-199a and miR-199b), and their variants, are not removed from the analysis. Once removed ambiguous sequences, the next filtering step was aimed to remove sequences that are likely to be sequencing errors. This second filtering step, referred to as relative abundance filter, occurred after the transformation of raw reads counts in reads per million (RPM), to account for differences in library size. Read counts were converted in RPM using the *cpm* function from the EdgeR package (version 3.40.2)[[9]](https://sciwheel.com/work/citation?ids=673952&pre=&suf=&sa=0&dbf=0). RPM for each miRNA (resulting from the canonical sequence and its isomiR sequences) were summed up, and the contribution of each sequence to the total miRNA’s RPM was estimated. Sequences whose average contribution across samples to total miRNA expression was < 1% were removed from the analysis. Concomitantly with the relative abundance filtering, also a low counts filtering was implemented to remove sequences with low expression (<3 counts in > 1 sample in EndoC-βH1; <3 counts in >6 samples in HI).

Filtered read counts were normalized with DESeq2’s (version 1.38.3)[[10]](https://sciwheel.com/work/citation?ids=129353&pre=&suf=&sa=0&dbf=0) median of the ratios method to account for differences in library size and RNA composition. Also the normalization process occurred independently for EndoC-βH1 and HI.

IsomiRs were classified according to their isoLabel based on the sRNAbench classification. In details, 8 different classes (or types) of miRNAs were identified:

- Canonical: if the read perfectly corresponds to the reference sequence on miRbase v22.1.
- Trim 3p: if the read matches the 5’end of the reference sequence but it has a shorter 3’ end.
- Ext 3p: if the read matches the 5’end of the sequence but it has a longer 3’ end.
- Trim 5p: if the read matches the 3’end of the reference but it has a shorter 5’ end.
- Ext 5p: if the read matches the 3’end of the reference sequence but it has a longer 5’ end.
- NTA: if the read has additional nucleotides at the 3’ end that do not match the precursor sequence.
- MultilVariant: if the read does not match neither the 3’ end and the 5’ end of the reference sequence. Also, isomiRs with composite modifications (extension and NTA) at 3’ end and/or 5’ end are included in this class.
- NucVar: the read has the same extremities of the reference sequence, but it shows a sequence variation.

Being isomiRs classification based on a hierarchy, sequences with different modifications can be assigned to the same class (i.e., sequences with both trimming and nucleotide variation will be assigned to the trimming class). For this reason, to detect isomiRs with a different seed sequence compared to the canonical counterpart, a seed conservation analysis was performed. For each isomiR, nucleotides in position 2-7 [[11]](https://sciwheel.com/work/citation?ids=1332638&pre=&suf=&sa=0&dbf=0) were identified as the seed sequence. The same procedure was performed for canonical miRNAs, according to miRbase reference sequence. Then, isomiRs’ seed sequences were compared to the seeds of their canonical counterparts. If differences in the sequences were detected, the isomiR was determined as ‘Not-conserved seed’ isomiR.

**In-silico Sequencing simulation**

The in-silico simulation was performed using a string of 22 elements (*synth_seq*), representing the canonical miRNA. The sequencing of the *synth_seq* was simulated with a probability of error in the reading of each nucleotide equal to 0,1% (Q=30). To model the error in the reading of each nucleotide with Q=30, a vector of *n=1000* elements, with *n=999* equal to False and *n=1* equal to True was defined (*error_vect*). Thus, the probability to sample an element from *error_vect* equal to True (1/1000) represents the probability of an error in the sequencing of the *synth_seq*.

The sequencing was simulated with the following procedure:

(i) For each nucleotide of the *synth_seq*, an element of *error_vect* was randomly sampled. (ii) If the sampled element was equal to *False*, the corresponding nucleotide of the sequencing simulation was set as equal to the nucleotide of the *synth_seq*, otherwise it was randomly selected a different nucleotide. This step resulted in the creation of a simulated read (referred to as *synth_read)*, which could be equal or not to the original *synth_seq.*

This 2-steps procedure was repeated n=*1.000.000* times, thus originating an equal number of *synth_reads* (referred to as *synth_pool*).

This pool of *synth_reads* was used for the validation of the relative abundance threshold. First, the occurrences (counts) and the relative abundance of each sequence from the *synth_pool* were estimated. Only sequences with 1 single sequencing error were used to model the probability distribution of false isomiRs. Indeed, sequences with more sequencing errors have few counts and, consequently, a very low relative abundance. This is because (i) there is a very low chance of having multiple sequencing errors and (ii) there is a high number of possible combinations. For this reason, a threshold on the relative abundance capable to remove false isomiRs with 1 single error will certainly remove sequences with multiple errors. The false isomiRs relative abundance distribution was estimated using a normal distribution. The goodness of the relative abundance threshold was computed as the probability of having an isomiR with a relative abundance >1%, given the previously estimated false isomiRs relative abundance distribution.

Among the reads originated from the sequencing’ simulation, n=*978.011* were equal to *synth_seq*.

On the other hand, the *21.989* reads detected as false isomiRs were assigned to 274 different sequences. Among them, *21.773* false isomiR reads (>99%) belonged to the *66* sequences with a single nucleotide variation compared to the *synth_seq*. The counts assigned to the single nucleotide variation sequences ranged from *298* to *377* counts. The relative abundance of these sequences to the total miRNA counts was estimated as the counts assigned to the sequence divided by the total number of reads (*1.000.000*). The Shapiro test assessed the normality of the distribution of their relative abundance (*p=0.23*). Thus, the probability distribution of false isomiRs was modelled with a normal distribution with the mean and standard deviation of their relative abundance (mean=*0.000330*, sd= *0,000017*). The probability of false isomiRs with a relative abundance higher than the threshold (*0.01*) was estimated with the *pnorm* function (*p=0*). The simulation corroborates the importance of the implementation of a filtering step based on the contribution to the corresponding miRNA counts. Indeed, false isomiRs with *>300* read counts were detected. These false isomiRs are likely to be kept from the low counts filtering. Thus, miRNAs with very high levels of expression (i.e., miR-16-5p) could give rise to sequencing errors that will be kept in the further analyses if no filters based on relative abundance are implemented.

**Statistical Analysis**

Regression analyses between isomiRs expression and clinical parameters were performed using multiple linear regression models. Normalized read counts were transformed in log2 scale, after the addition of a pseudo count. The regressions were modelled using the log2-transformed isomiR expression as dependent variable and the clinical parameter, the age, the gender and the BMI as regressors. To exclude influent values from the regression analysis, the cooks’ distance of each point for each model was estimated. Points with a cooks’ distance > 5 folds the average cooks’ distance of the points in the regression were removed as influent points. After this step that avoids regressions strongly driven from a single data point, linear models were estimated for each isomiR with each clinical parameter. Regressions with a p-value associated to the coefficient assigned to the clinical parameter *< 0.05* were considered as statistically significant, independently from the effect of the covariates (age, gender, and BMI). The *partial* *R2* of the clinical parameter was estimated using the function *partial_r2* from the sensemakr package [[12]](https://sciwheel.com/work/citation?ids=15613047&pre=&suf=&sa=0&dbf=0). The square root of the *partial R2* was then estimated and multiplied by the sign of the coefficient, thus originating the *partial R*, that is also informative about the sign of the regression. For representative purposes, the log2-transformed counts were corrected for the effect of the covariates for each regression. The correction was performed by estimating the effect of the covariates in each regression as the coefficient assigned to the covariate multiplied by the value of the covariate. Then, the effect of the covariates for each data point in the regression was summed up and then subtracted to the normalized log2-scaled counts. With this procedure was estimated, and then removed, the overall effect of the covariates on the dependent variable (isomiR expression).

Hierarchical clustering analysis of miRNAs composition was performed with the pheatmap package (Version 1.0.12) [[13]](https://sciwheel.com/work/citation?ids=15613117&pre=&suf=&sa=0&dbf=0). The composition of each miRNA was computed as the percentage of the counts assigned to the previously defined miRNA classes, averaged across samples. The dissimilarity matrix was estimated using the Euclidean distance, while the complete linkage method was used for the agglomeration. The optimal number of clusters was determined using the silhouette method (NBclust package Version 3.0.1) [[14]](https://sciwheel.com/work/citation?ids=15613118&pre=&suf=&sa=0&dbf=0), with a number of clusters ranging from 2 to 10. The number of clusters that maximize the average silhouette was determined as the optimal number of clusters. Similarity in miRNAs composition among clusters detected in HI and EndoC-βH1 was computed using the Jaccard similarity coefficient. The similarity analysis was restricted to miRNAs commonly detected in the two experimental groups (n=140 miRNAs). The Jaccard similarity index was computed as the intersection above the union of the miRNAs detected for the different pairs of clusters.

To identify isomiR sequences with major contribution to the experimental group, a delta ranking analysis was performed. For each experimental group (HI and EndoC-βH1) sequences were ranked based on their expression in a descending way (from high to low expression). Thus, sequences with high expression have a low rank and vice versa. Then, the difference in rank (delta rank) between the isomiR and its corresponding canonical counterpart was computed. Highly negative values of delta rank are related to isomiR sequences whose reference sequence has a higher rank and, consequently, a lower expression. The analysis was restricted to the n=50 most expressed sequences to avoid the identification of high delta ranking related to sequences with low expression.

**IsomiRdb analysis**

IsomiRdb datasets were downloaded and re-analysed to validate the isomiR signature in beta cells and other cell types. Sample metadata, miRNA expression (Read Per Million), and isomiR expression (Reads Per Million) files were retrieved form IsomiRdb and re-analysed using a multi-step filtering.

Firstly, metadata information was used to select sequencing data derived from healthy and primary cell studies. At first, only sources derived from Sequence Read Archive (SRA) [[15]](https://sciwheel.com/work/citation?ids=2084729&pre=&suf=&sa=0&dbf=0) and in which cell origin is present were retained. From the remaining dataset, external IDs were used to collect study title and attributes from NCBI (https://www.ncbi.nlm.nih.gov/). On the latter were applied additional filters to produce the final metadata contain 396 sequencing derived from 99 cell types. At this stage, IsomiR expression sequencing were filtered using previous mentioned sample metadata, and the mean expression of isomiRs for each cell was estimated. The same procedure was also applied to the miRNA expression file. Therefore, isomiRs and miRNA dataframes were merged to apply relative abundance filter. IsomiRs with a relative abundance averaged across all cell types > 1% were kept. From the resulting dataframe, Z-score were calculated as follow:

$$Z= \frac{x_{i}-\mu}{\sigma}$$

Where $x_{i}$ is the mean expression of a given isomiR, $\mu$ is the mean expression in all cells, and $\sigma$ the standard deviation.

**ESM TABLES**

**ESM Table 1.** Demographic and clinical-metabolic characteristics of non-diabetic living donors.

| **Case ID** | **Gender (M/F)** | **Age (y)** | **BMI (kg/m^2^)** | **Basal Glucose (mmol/l)** | **Mean Glucose (mmol/l)** | **Basal Insulin (pmol/l)** | **Mean Insulin (pmol/l)** | **Basal Insulin Secretion Rate, ISR (pmol.min-1.m-2)** | **Total Insulin Secretion Rate, ISR (pmol.min-1.m-2)** | **Glucose Sensitivity (pmol min^-1^m^-2^mM^-1^)** | **Rate Sensitivity (nmol m^-2^mM^-1^)** |
| --- | --- | --- | --- | --- | --- | --- | --- | --- | --- | --- | --- |
| LCM-4 | M | 71 | 28,60 | 5,11 | 9,30 | 42,60 | 329,18 | 71,10 | 56,18 | 83,25 | 430,31 |
| LCM-5 | M | 79 | 30,12 | 5,77 | 9,51 | 30,00 | 447,98 | 69,09 | 47,27 | 75,75 | 192,75 |
| LCM-6 | F | 76 | 24,09 | 4,44 | 6,91 | 31,20 | 334,43 | 47,94 | 39,38 | 92,41 | 997,47 |
| LCM-7 | F | 56 | 21,70 | 4,33 | 7,45 | 24,60 | 174,90 | 45,72 | 30,49 | 64,78 | 328,65 |
| LCM-8 | F | 67 | 31,43 | 4,83 | 7,47 | 67,20 | 376,05 | 90,97 | 50,61 | 116,95 | 0,00 |
| LCM-9 | M | 78 | 29,10 | 4,88 | 7,22 | 11,40 | 96,15 | 44,48 | 16,28 | 26,56 | 656,03 |
| LCM-10 | M | 66 | 28,10 | 6,33 | 8,61 | 73,20 | 465,08 | 109,35 | 52,03 | 65,18 | 966,09 |
| LCM-11 | F | 58 | 28,39 | 5,27 | 7,56 | 69,60 | 295,73 | 61,04 | 33,49 | 101,90 | 0,00 |
| LCM-12 | M | 72 | 21,63 | 5,16 | 8,77 | 30,60 | 281,40 | 48,51 | 35,43 | 49,83 | 337,64 |
| LCM-13 | M | 58 | 24,22 | 5,05 | 7,38 | 42,00 | 291,75 | 52,44 | 41,70 | 91,60 | 2766,54 |
| LCM-14 | F | 45 | 25,14 | 5,22 | 6,02 | 27,60 | 88,88 | 38,64 | 25,54 | 74,72 | 276,47 |
| LCM-15 | F | 66 | 22,04 | 5,38 | 6,69 | 43,80 | 444,75 | 60,61 | 44,78 | 113,39 | 1136,28 |
| LCM-16 | M | 62 | 21,15 | 4,94 | 6,14 | 26,40 | 178,50 | 36,79 | 25,34 | 61,05 | 2076,63 |
| LCM-17 | F | 51 | 22,15 | 4,88 | 7,60 | 48,00 | 420,98 | 34,89 | 42,97 | 100,21 | 329,25 |
| LCM-18 | M | 59 | 19,93 | 4,00 | 7,55 | 13,20 | 174,30 | 53,56 | 50,00 | 94,81 | 778,06 |
| LCM-19 | F | 78 | 24,27 | 5,33 | 7,92 | 49,80 | 428,93 | 96,77 | 83,31 | 189,36 | 2953,03 |

**ESM Table 2.** Characteristics of non-diabetic multiorgan donors recruited wihtin INNODIA EUnPOD network.

| **Case ID** | **Gender (M/F)** | **Age (y)** | **BMI (kg/m^2^)** | **AutoAb (ELISA)** | **HiRes HLA** | **Cause of death** |
| --- | --- | --- | --- | --- | --- | --- |
| LCM-1 | M | 39 | 23,6 | GADA negative, IA-2A negative, ZnT8A negative | HLA:A*03,33; B*14, B*14 SD B65; C*08; DRB1*01, DQB1*05 | Trauma |
| LCM-2 | M | 49 | 25,8 | GADA negative, IA-2A negative, ZnT8A negative | HLA:A*03,68; B*35,47; C*04,06; DRB1*03,08; DQB1*02,04 | Cardiovascular disease |
| LCM-3 | F | 46 | 32,5 | GADA negative, IA-2A negative, ZnT8A negative | HLA:A*24; B*15,18; C*7; DRB1*04,11; DQB1*3 | Cardiovascular disease |

**ESM Table 3.** Table showing RNA integrity Number (RIN, scale 0-10) and concentration (pg/µl) of total RNA extracted from LCM islets captured from pancreatic tissue sections of non-diabetic subjects and EndoC-βH1 samples.

| **Case ID** | **RNA Integrity Number (RIN) value** | **Concentration (pg/µl)** |
| --- | --- | --- |
| LCM-1 | 6,1 | 905 |
| LCM-2 | 6 | 271 |
| LCM-3 | 7,9 | 285 |
| LCM-4 | 5,9 | 326 |
| LCM-5 | 6 | 292 |
| LCM-6 | 5,8 | 348 |
| LCM-7 | 7,5 | 195 |
| LCM-8 | 6,3 | 990 |
| LCM-9 | 6,5 | 231 |
| LCM-10 | 7,1 | 489 |
| LCM-11 | 5,6 | 369 |
| LCM-12 | 7,3 | 193 |
| LCM-13 | 7,2 | 599 |
| LCM-14 | 6 | 391 |
| LCM-15 | 5,9 | 310 |
| LCM-16 | 7,2 | 236 |
| LCM-17 | 6,8 | 349 |
| LCM-18 | 7,5 | 660 |
| LCM-19 | 5,2 | 1085 |
| ENDOC-1 | 8,5 | 995 |
| ENDOC-2 | 9,3 | 1000 |
| ENDOC-3 | 8,5 | 1000 |

**ESM Table 4.** Table showing the Jaccard similarity index across the different pairs of Clusters detected in HI and EndoC-βH1 samples.

| **Clusters** | HI Cluster Ext 3p | HI Cluster MultiVariant | HI Cluster NTA | HI Cluster Trim 3p | HI Cluster Trim 5p | HI Cluster Canonical |
| --- | --- | --- | --- | --- | --- | --- |
| EndoC-βH1 Cluster Canonical | 0,01 | 0,00 | 0,03 | 0,01 | 0,00 | 0,78 |
| EndoC-βH1 Cluster Ext 3p | 0,61 | 0,04 | 0,00 | 0,00 | 0,00 | 0,09 |
| EndoC-βH1 Cluster MultiVariant | 0,00 | 0,20 | 0,00 | 0,03 | 0,00 | 0,02 |
| EndoC-βH1 Cluster NTA | 0,00 | 0,00 | 0,63 | 0,00 | 0,00 | 0,00 |
| EndoC-βH1 Cluster Trim 3p | 0,00 | 0,00 | 0,03 | 0,79 | 0,00 | 0,04 |
| EndoC-βH1 Cluster Trim 5p | 0,00 | 0,00 | 0,00 | 0,00 | 1,00 | 0,00 |

**ESM Table 5.** Table showing the characteristics of the beta cell isomiRs signature. The isomiR’s sequence, its mature name, the isomiR class and the preservation of the seed sequence are reported for each isomiR.

| **Sequence** | **miRNA** | **isomiR Type** | **Seed** |
| --- | --- | --- | --- |
| TTCCCTTTGTCATCCTATGCCTG | hsa-miR-204-5p | Ext 3p 1 | Conserved |
| TAATACTGCCGGGTAATGATGG | hsa-miR-200c-3p | Trim 3p -1 | Conserved |
| TGGCTCAGTTCAGCAGGAACAGT | hsa-miR-24-3p | NTA | Conserved |
| TTCAAGTAATCCAGGATAGG | hsa-miR-26a-5p | Trim 3p -2 | Conserved |
| TCAAGAGCAATAACGAAAAATG | hsa-miR-335-5p | Trim 3p -1 | Conserved |
| GTGAGGACTCGGGAGGTGGAGGGT | hsa-miR-1224-5p | Ext 3p 5 | Conserved |
| AGATCGACCGTGTTATATTCG | hsa-miR-369-5p | Trim 3p -1 | Conserved |
| GTGAGGACTCGGGAGGTGGAG | hsa-miR-1224-5p | Ext 3p 2 | Conserved |
| TGTAAACATCCTTGACTGGAAGCT | hsa-miR-30e-5p | Ext 3p 2 | Conserved |
| GTGGGGGAGAGGCTGT | hsa-miR-1275 | Trim 3p -1 | Conserved |
| GCCTGCTGGGGTGGAACCTGG | hsa-miR-370-3p | Trim 3p -1 | Conserved |
| AGCTACATTGTCTGCTGGGTTT | hsa-miR-221-3p | Trim 3p -1 | Conserved |
| TTCAAGTAATTCAGGATAGGTT | hsa-miR-26b-5p | Ext 3p 1 | Conserved |
| TACTGCATCAGGAACTGATTGG | hsa-miR-217-5p | Trim 3p -1 | Conserved |
| AGAGGTAGTAGGTTGCATAGT | hsa-let-7d-5p | Trim 3p -1 | Conserved |
| TAATCTCAGCTGGCAACTGTG | hsa-miR-216a-5p | Trim 3p -1 | Conserved |
| TGGTAGACTATGGAACGTAG | hsa-miR-379-5p | Trim 3p -1 | Conserved |
| TGTAAACATCCCCGACTGGAAGCT | hsa-miR-30d-5p | Ext 3p 2 | Conserved |
| TCCCTGAGACCCTTTAACCTGTG | hsa-miR-125a-5p | Trim 3p -1 | Conserved |
| GTGAGGACTCGGGAGGTGGAGGG | hsa-miR-1224-5p | Ext 3p 4 | Conserved |
| AAGCATTCTTTCATTGGTTGGT | hsa-miR-1179 | Ext 3p 1 | Conserved |
| TGGAAGACTAGTGATTTTGTTGT | hsa-miR-7-5p | Trim 3p -1 | Conserved |
| AGAGGCTGGCCGTGATGAATTCG | hsa-miR-485-5p | Ext 3p 1 | Conserved |
| ATAGTAGACCGTATAGCGTACG | hsa-miR-411-5p | Ext 5p 1 | NotConserved |
| TGTAACAGCAACTCCATGTGG | hsa-miR-194-5p | Trim 3p -1 | Conserved |
| GTGAGGACTCGGGAGGTGGA | hsa-miR-1224-5p | Ext 3p 1 | Conserved |
| TAATACTGCCTGGTAATGATG | hsa-miR-200b-3p | Trim 3p -1 | Conserved |
| TCCTGTACTGAGCTGCCCCGAGT | hsa-miR-486-5p | NTA | Conserved |
| GTGAGGACTCGGGAGGTGGAGG | hsa-miR-1224-5p | Ext 3p 3 | Conserved |
| TCTTGGAGTAGGTCATTGGGTGT | hsa-miR-432-5p | NTA | Conserved |
| TGCGGGGCTAGGGCTAACAGC | hsa-miR-744-5p | Trim 3p -1 | Conserved |
| TGACCTATGAATTGACAGCCAGT | hsa-miR-192-5p | MultiVariant | NotConserved |
| TAGCTTATCAGACTGATGTTG | hsa-miR-21-5p | Trim 3p -1 | Conserved |
| TGTAAACATCCCCGACTGGAAGC | hsa-miR-30d-5p | Ext 3p 1 | Conserved |
| TAGCACCATTTGAAATCGGTT | hsa-miR-29c-3p | Trim 3p -1 | Conserved |
| ATCACATTGCCAGGGATTACC | hsa-miR-23b-3p | Trim 3p -2 | Conserved |
| TCCCTGAGACCCTTTAACCTGTGT | hsa-miR-125a-5p | NTA | Conserved |
| ACTCGGCGTGGCGTCGGTCGTGG | hsa-miR-1307-3p | Ext 3p 1 | Conserved |
| TTCAACGGGTATTTATTGAGC | hsa-miR-95-3p | Trim 3p -1 | Conserved |
| TAACACTGTCTGGTAACGATGTT | hsa-miR-200a-3p | Ext 3p 1 | Conserved |
| TAACAGTCTACAGCCATGGTCGT | hsa-miR-132-3p | NTA | Conserved |
| TAACACTGTCTGGTAAAGATG | hsa-miR-141-3p | Trim 3p -1 | Conserved |
| TCTTGGAGTAGGTCATTGGGT | hsa-miR-432-5p | Trim 3p -2 | Conserved |
| TACCCTGTAGAACCGAATTTGT | hsa-miR-10b-5p | Trim 3p -1 | Conserved |
| CGAATGTTGCTCGGTGAACCCCT | hsa-miR-409-3p | Ext 5p 1 | NotConserved |
| TCCCTGAGACCCTTTAACCTGT | hsa-miR-125a-5p | Trim 3p -2 | Conserved |

**ESM FIGURES**

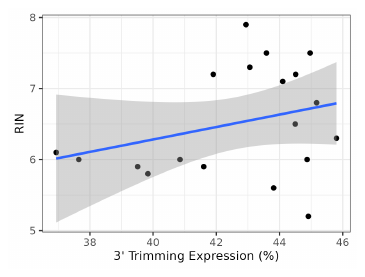

**ESM Figure 1**. Correlation between 3’ Trimming Expression and RNA Integrity Number (RIN) in LCM Human pancreatic islets. The scatterplot shows the association between the RIN (from 0-10) and the proportion (reported as percentage) of expression assigned to 3’ Trim isomiRs. Each dot represents a LCM-HI samples. Regression line (in blue) and intervals (grey shadowed area) are reported.

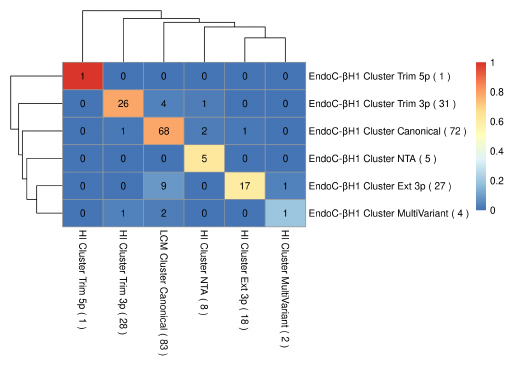

**ESM Figure 2.** Jaccard matrix showing miRNAs clusters concordance between HI and EndoC-βH1. The concordance between clusters was computed using the Jaccard similarity coefficient, and the analysis was restricted to the annotated miRNAs commonly detected in the two experimental groups. The colour represents the Jaccard similarity coefficient, while the values reported represent the intersection of the miRNAs detected in the pairs of clusters. The values reported in brackets alongside the name of the cluster represent the number of miRNAs present in the cluster.

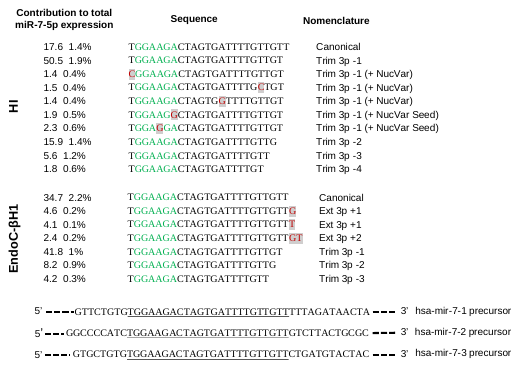

**ESM Figure 3**. Scheme showing the different sequences assigned to miR-7-5p in HI and EndoC-βH1. In the scheme is reported the average percentage (with standard deviation) of the contribution of the sequence to the total expression of miR-7-5p. In the bottom are represented the three different precursor sequences of the miRNA. Both in HI and EndoC-βH1 the most expressed sequence derives from a single nucleotide trimming at the 3’ end. Moreover, in HI, 2 out of the 6 sequences with a single nucleotide trimming have also a NucVar in the seed sequence, potentially targeting different genes compared to the canonical miRNA.

**ESM REFERENCES**

[1.    Mezza T, Clemente G, Sorice GP, et al (2015) Metabolic consequences of the occlusion of the main pancreatic duct with acrylic glue after pancreaticoduodenectomy. Am J Surg 210:783–789. https://doi.org/10.1016/j.amjsurg.2014.12.052](https://sciwheel.com/work/bibliography/15612984)

[2.    Brusco N, Sebastiani G, Di Giuseppe G, et al (2023) Intra-islet insulin synthesis defects are associated with endoplasmic reticulum stress and loss of beta cell identity in human diabetes. Diabetologia 66:354–366. https://doi.org/10.1007/s00125-022-05814-2](https://sciwheel.com/work/bibliography/13977332)

[3.    American Diabetes Association Professional Practice Committee (2022) 2. Classification and Diagnosis of Diabetes: Standards of Medical Care in Diabetes-2022. Diabetes Care 45:S17–S38. https://doi.org/10.2337/dc22-S002](https://sciwheel.com/work/bibliography/12466885)

[4.    Mezza T, Ferraro PM, Di Giuseppe G, et al (2021) Pancreaticoduodenectomy model demonstrates a fundamental role of dysfunctional β cells in predicting diabetes. J Clin Invest 131:. https://doi.org/10.1172/JCI146788](https://sciwheel.com/work/bibliography/11524219)

[5.    Mari A, Tura A, Natali A, et al (2010) Impaired beta cell glucose sensitivity rather than inadequate compensation for insulin resistance is the dominant defect in glucose intolerance. Diabetologia 53:749–756. https://doi.org/10.1007/s00125-009-1647-6](https://sciwheel.com/work/bibliography/9771328)

[6.    Ravassard P, Hazhouz Y, Pechberty S, et al (2011) A genetically engineered human pancreatic β cell line exhibiting glucose-inducible insulin secretion. J Clin Invest 121:3589–3597. https://doi.org/10.1172/JCI58447](https://sciwheel.com/work/bibliography/1914391)

[7.    Grieco GE, Sebastiani G, Fignani D, et al (2021) Protocol to analyze circulating small non-coding RNAs by high-throughput RNA sequencing from human plasma samples. STAR Protocols 2:100606. https://doi.org/10.1016/j.xpro.2021.100606](https://sciwheel.com/work/bibliography/11247261)

[8.    Aparicio-Puerta E, Gómez-Martín C, Giannoukakos S, et al (2022) sRNAbench and sRNAtoolbox 2022 update: accurate miRNA and sncRNA profiling for model and non-model organisms. Nucleic Acids Res 50:W710–W717. https://doi.org/10.1093/nar/gkac363](https://sciwheel.com/work/bibliography/15387713)

[9.    Robinson MD, McCarthy DJ, Smyth GK (2010) edgeR: a Bioconductor package for differential expression analysis of digital gene expression data. Bioinformatics 26:139–140. https://doi.org/10.1093/bioinformatics/btp616](https://sciwheel.com/work/bibliography/673952)

[10.   Love MI, Huber W, Anders S (2014) Moderated estimation of fold change and dispersion for RNA-seq data with DESeq2. Genome Biol 15:550. https://doi.org/10.1186/s13059-014-0550-8](https://sciwheel.com/work/bibliography/129353)

[11.   Agarwal V, Bell GW, Nam J-W, Bartel DP (2015) Predicting effective microRNA target sites in mammalian mRNAs. eLife 4:. https://doi.org/10.7554/eLife.05005](https://sciwheel.com/work/bibliography/1332638)

[12.   Cinelli C, Ferwerda J, Hazlett C (2021) sensemakr: Sensitivity Analysis Tools for Regression Models](https://sciwheel.com/work/bibliography/15613047)

[13.   Kolde R (2019) pheatmap: Pretty Heatmaps](https://sciwheel.com/work/bibliography/15613117)

[14.   Charrad M, Ghazzali N, Boiteau V, Niknafs A (2014) NbClust: An R Package for Determining the Relevant Number of Clusters in a Data Set. Journal of Statistical Software 61:1–36](https://sciwheel.com/work/bibliography/15613118)

[15.   Leinonen R, Sugawara H, Shumway M, International Nucleotide Sequence Database Collaboration (2011) The sequence read archive. Nucleic Acids Res 39:D19-21. https://doi.org/10.1093/nar/gkq1019](https://sciwheel.com/work/bibliography/2084729)
